# Within-host competition sparks pathogen molecular evolution and perpetual microbiota dysbiosis

**DOI:** 10.1101/2024.09.03.610829

**Authors:** E.J. Stevens, J.D. Li, T.E. Hector, G.C. Drew, K. Hoang, S.T.E. Greenrod, S. Paterson, K.C. King

**Affiliations:** Department of Biology, University of Oxford, Oxford, UK; School of Life Sciences, Keele University, Keele, UK; Division of Infectious Diseases, Emory University School of Medicine, Atlanta, US; Institute of Infection, Veterinary and Ecological Sciences, University of Liverpool, Liverpool, UK; Department of Zoology, University of British Columbia, Vancouver, Canada; Department of Microbiology & Immunology, University of British Columbia, Vancouver, Canada

## Abstract

Pathogens newly invading a host must compete with resident microbiota. This within-host microbial warfare could lead to more severe disease outcomes or constrain the evolution of virulence. Using experimental evolution of a widespread pathogen (*Staphylococcus aureus*) and a native microbiota community in *C. elegans* nematode hosts, we show that a competitively superior pathogen displaced microbiota and reduced species richness, whilst maintaining virulence across generations. Conversely, pathogen populations and microbiota passaged separately caused more host harm relative to their respective ancestral controls. We find the evolved increase in virulence exhibited by pathogen populations passaged independently (compared to ancestral controls) was partly mediated by enhanced expression of the global virulence regulator *agr* and increased biofilm formation. Whole genome sequencing revealed shifts in the mode of selection from directional (on pathogens evolving alone) to fluctuating (on pathogens evolving with a host microbiota), with competitive interactions driving early diversification among pathogen populations. Metagenome sequencing of the evolved microbiota shows that evolution in infected hosts caused a significant reduction in community stability, along with restrictions on the co- existence of some species based on nutrient competition. Our study reveals how microbial competition during emerging infection determines the patterns and processes of evolution with major consequences for host health.

## Introduction

During host invasion, pathogens encounter a microbial community (i.e. microbiota) occupying the intended niche. Unlike resident microbiota, invading pathogens are not locally adapted. This lack of evolutionary history with the host may put invading pathogens at a competitive disadvantage. Indeed, competitive exclusion of pathogens, and thus host protection, by microbiota is a common phenomenon observed across a range of animal hosts ^1,2^. To overcome this ecological challenge, pathogens have been shown to kill competitors via toxins (i.e., interference competition) or provoke host inflammation in a type of ‘proactive invasion’ (reviewed in^3,4^. Pathogens can also compete with microbiota for host resources ^5–7^. These strategies might push out resident microbiota, reducing their numbers and diversity ^4,8,9^, and contribute to high pathogen virulence during acute infection ^10–12^. The potential for competitive interactions between emerging pathogens and host microbiota to affect disease severity is unclear, but has generated considerable interest for managing the harm caused by infection in human medicine ^13,14^, wildlife conservation ^15^, and agriculture ^16^.

Within-host microbial warfare can thus come with huge fitness implications for hosts, microbiota, and pathogens ^17^. The evolutionary outcomes of microbiota-pathogen competition on disease severity are complex. Higher pathogen virulence might evolve over time in a more competitively exclusive microbiome ^17^, or be limited if already at a high optimum ^18^. However, pathogens should be favoured to exploit their hosts cautiously to avoid killing them prematurely ^19^. Microbiota of healthy hosts can exhibit rapid evolutionary dynamics in response to aging and diet ^20^, with resource competition as a strong source of selection ^21–24^. In the presence of competition with pathogens, individual microbial commensal species can become more competitive ^7^ and/or protective ^25^. Alternatively, resource competition might limit microbial evolutionary responses ^26,27^. Within-host evolutionary interactions between pathogens and microbiota may be an important process underlying the transition from commensalism to pathogenicity ^28^.

Here, we directly tested whether pathogen competition with host microbiota could shape the evolution of disease severity. We experimentally evolved a widespread, disease-causing animal pathogen (*Staphylococcus aureus*) using a *C. elegans* nematode worm model of infection. *C. elegans* are likely exposed to *Staphylococcus* spp. in natural environments ^29–31^.

However, in our model, *S. aureus* acts as a novel invading pathogen to canonical laboratory *C. elegans* nematodes. Pathogenic *S. aureus* strains are known to infect a diversity of animal host species, including domestic and wild animals ^29,30,32–34^, where they can engage in (in some cases toxin-mediated) resource competition with other microbes ^35–38^. Nematodes infected by *S. aureus* are harmed when the pathogen accumulates and produces toxins in the host intestine ^39–41^. To independently test whether microbiota could become more exclusionary during *S. aureus* infection, we also passaged a community of naturally-associated microbes ^42^ within nematode populations. Across short and longer-term time scales, we examined the eco-evolutionary trajectories of each passaged pathogen population and microbiota community in the context of within-host competition and virulence. The molecular and mechanistic basis of microbial adaptive processes were also explored.

## Results

### Presence of a microbiota in the *C. elegans* gut increases host mortality from *S. aureus* infection

We tested how disease severity from *S. aureus* was initially affected by host microbiota. When infected with *S. aureus,* we found mortality of *C. elegans* nematodes significantly increased in hosts pre-colonised by a microbiota community compared to *S. aureus* alone (fig. 1A). Colonisation by the microbiota alone did not yield any host mortality (fig. 1A). We determined whether this increased host-killing observed during infection was dependent on the whole microbiota community or one species. We exposed nematode populations to each microbiota species separately and infected them with *S. aureus*. We found considerable variation in the extent to which colonisation by each microbiota component contributed to host killing by the pathogen (fig. 1Ai). Significantly higher host mortality was observed in nematodes colonised with CEent1, JUb66 and BIGb393 during *S. aureus* infection, compared to the *E. coli* OP50 control (standard *C. elegans* food). Colonisation by strains MYb10, MYb71, BIGb0170 and MYb11 did not result in significantly higher mortality during *S. aureus* infection. In the absence of *S. aureus*, these species do not individually cause mortality in *C. elegans* ^42^.

**Figure 1.**
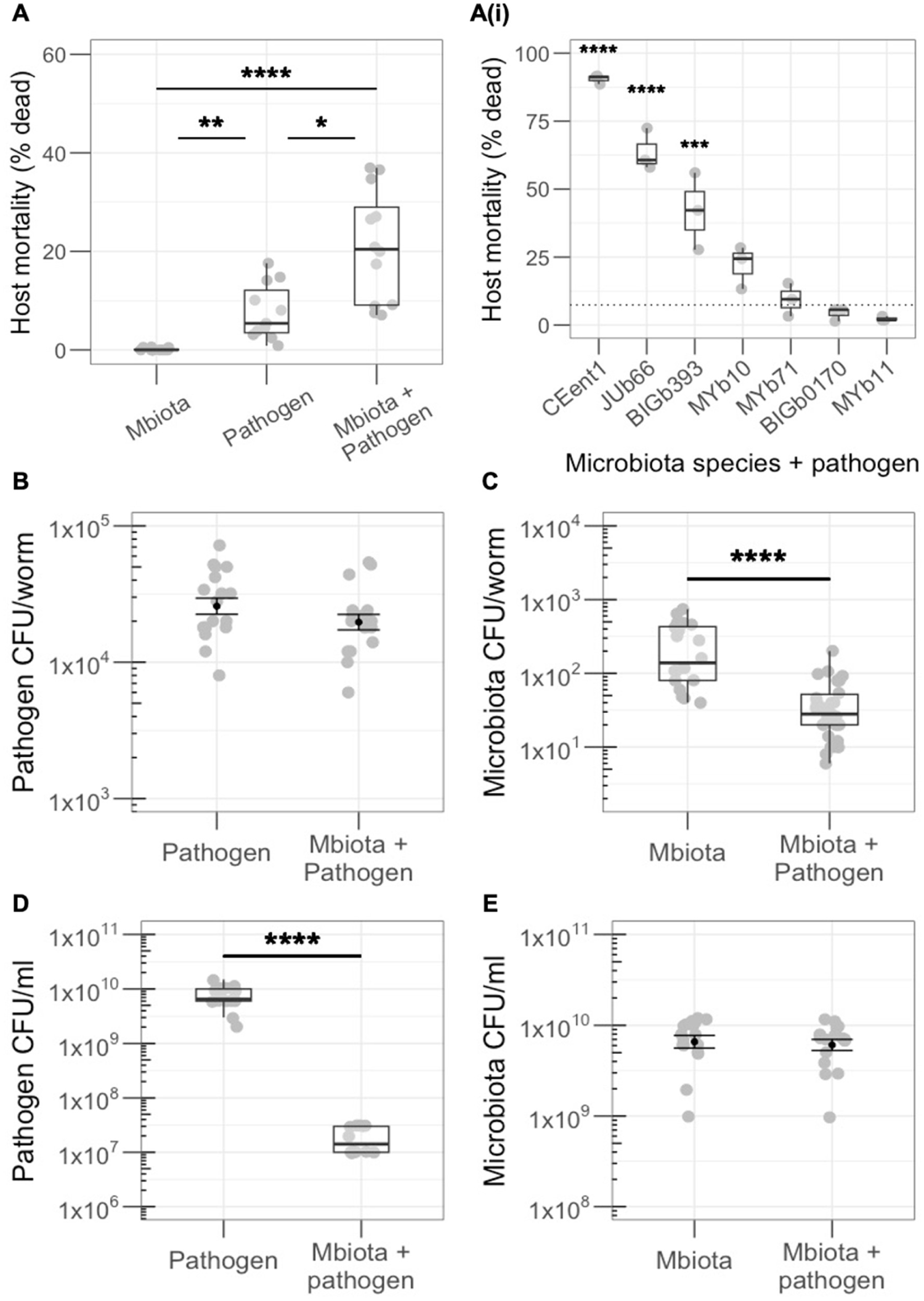
**A)** Mortality of *C. elegans* from *S. aureus* is significantly higher in the presence vs absence of a 7-species microbiota community (n=11-12, Kruskal-Wallis rank sum test, df = 2, p<0.0001, Dunn test comparisons: Pathogen vs Mbiota + pathogen p=0.053, Pathogen vs Mbiota p=0.003, Mbiota vs Mbiota + pathogen p<0.0001). Ai) Significant variation in the extent to which each microbiota component individually facilitates pathogen virulence in *C. elegans*. Significance is indicated only where species are significantly different to the control *E. coli* OP50, depicted by dotted line (n=3, linear regression, f=59.17, df=16, p<0.0001, pairwise comparisons using ‘emmeans’ package: CEent1 vs control p<0.0001, JUb66 vs control p<0.0001, BIGb393 vs control p=0.002). **B)** *In vivo* colonisation of *C. elegans* by the pathogen is not significantly affected by the presence of the microbiota (n=18, Welch Two-sample T test p=0.172), while **C)** colonisation by the microbiota is significantly lower in the presence vs absence of the pathogen (n=20-30, Wilcoxon rank sum test p<0.0001), as measured by colony forming units (CFU) per worm. **D)** *In vitro* growth of the pathogen is significantly reduced in the presence vs absence of the microbiota in LB media (n=12-18, Wilcoxon rank sum test p<0.0001), as measured by CFU/ml. **E)** *In vitro* growth of the microbiota is comparable in the presence and absence of the pathogen in LB media (n=17- 18, Welch Two-sample T test p=0.474).

We tested whether differences in colonisation of *C. elegans* by the pathogen and microbiota determined disease severity. We extracted internal bacteria from surface-sterilised nematodes and counted the number of colony forming units (CFU) per host. Pathogen load remained unchanged by the presence of host microbiota (fig 1B), thus there is no relationship between pathogen-induced host mortality and *in vivo* load. The microbiota is a weaker competitor within the host, as we found total microbiota load was significantly reduced during pathogen infection (fig 1C). We observed the opposite dynamic during *in vitro* competition, with pathogen growth suppressed in the presence of the microbiota (fig 1D), while growth of the microbiota was consistent in the presence and absence of *S. aureus* (fig 1E). Growth of, and competition dynamics between, the pathogen and microbiota are therefore dependent on the host environment.

### *S. aureus* becomes more virulent compared to controls when evolved in isolation

We hypothesised that competition with host microbiota would drive changes in pathogen virulence over evolutionary time. We passaged *S. aureus* either with a non-evolving (ancestral) 7-species microbiota or on its own, within genetically homogeneous, non-evolving *C. elegans* populations for 15 passages (fig 2A). The 7-species microbiota community was similarly passaged with ancestral *S. aureus*. No-host controls were included to act as a proxy for the ancestor, controlling for any lab adaptation that may have occurred during the experiment. Figure 2B depicts each of the evolving bacterial groups. All groups consisted of six replicate populations. Pathogen populations started from the same clone of *S. aureus*, and microbiota communities from the same clone of each species. Evolution was thus dependent on *de novo* mutation and follow-on selection. At the end of each passage, 100 colonies of *S. aureus* and 100 colonies from the mixed microbiota communities were collected randomly from nematodes and used to start the next passage.

**Figure 2.**
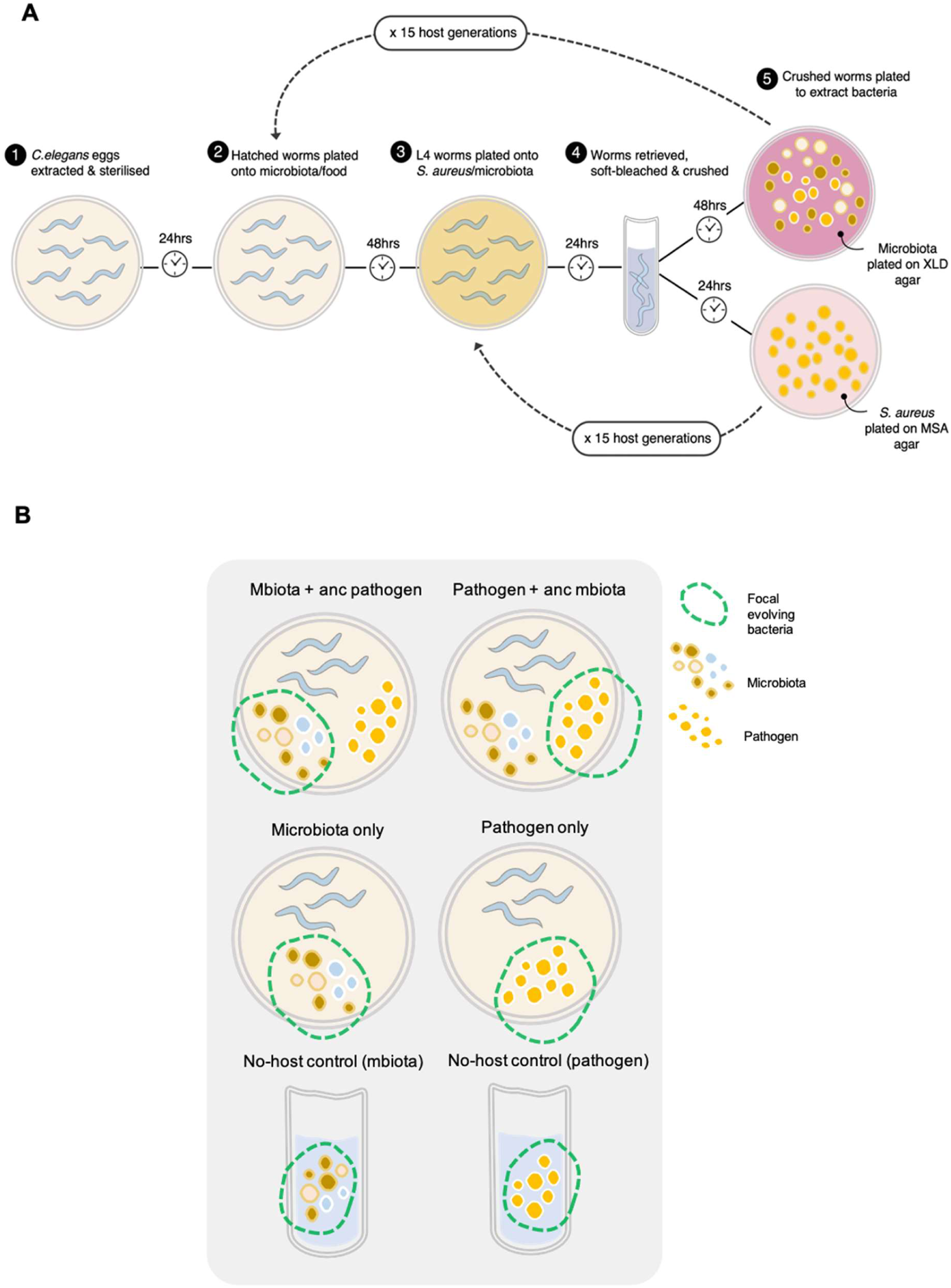
**A)** Overview of the experimental evolution method. **B)** Overview of each evolving group in the experiment.

**Figure 3.**
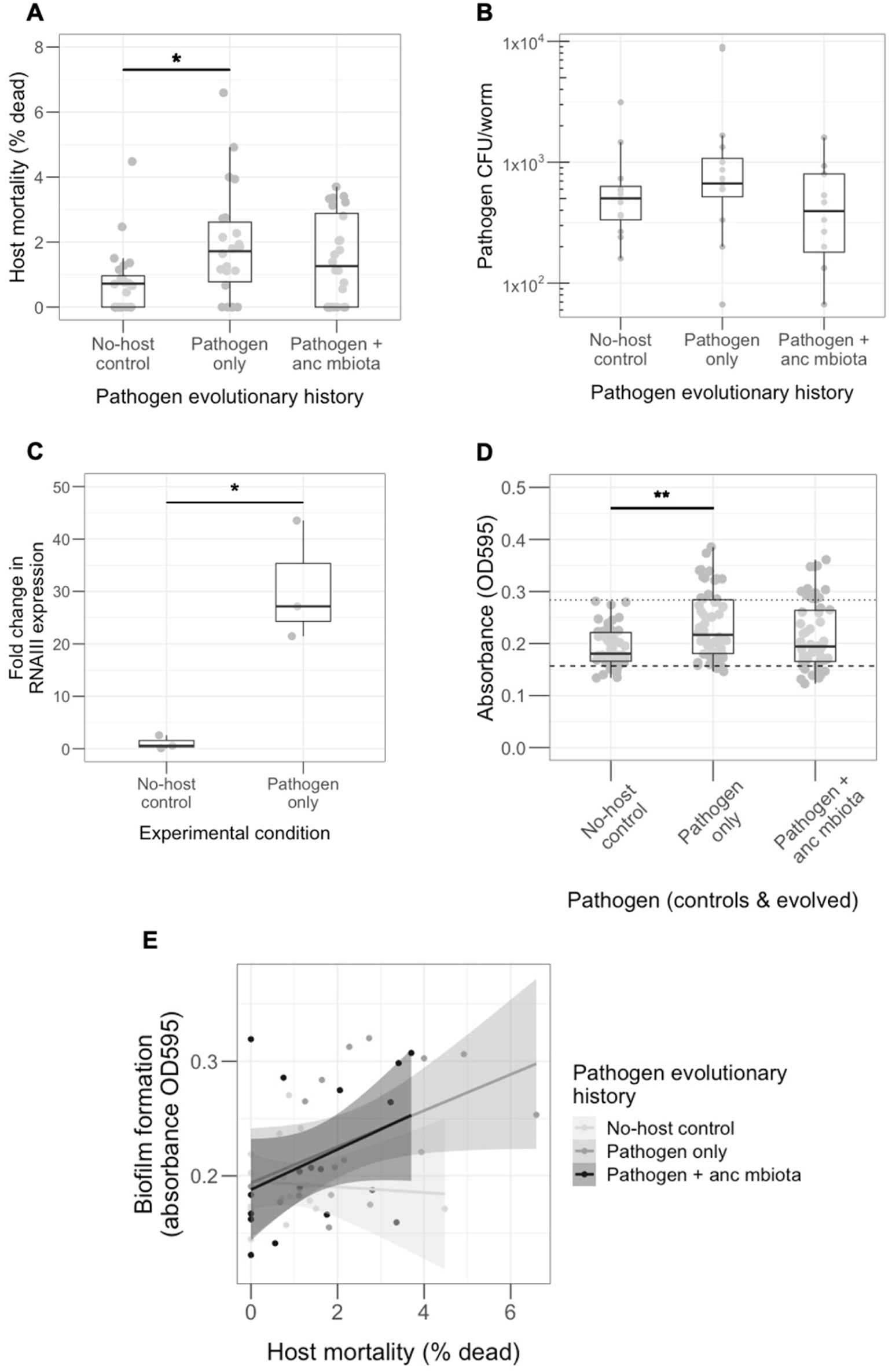
All data shown represents the 15th (final) evolved passage. **A)** The ‘Pathogen only’ group was significantly more virulent in the absence of the microbiota than the ‘No-host control’ (n=22-24, Kruskal Wallis rank sum test, df=2, p=0.038, Dunn test comparison No-host control vs Pathogen only p=0.038). **B)** The difference in virulence is not mediated by bacterial colonisation of *C. elegans*, which is comparable across all evolved pathogens (n=12-16, Kruskal Wallis rank sum test, df=2, p=0.216). **C)** Expression of the global virulence regulator *agr*, as measured by RNAIII expression, is significantly higher in the ‘Pathogen only’ group compared to the ‘No-host control’ (n=3, Welch Two-sample T test p=0.045). **D)** Evolved pathogens were assayed for their ability to form biofilm in a 96-well plate, alongside the poor biofilm-forming strain LAC (negative control – dashed line), and MSSA476 (positive control – dotted line). The ‘Pathogen only’ group showed significantly more biofilm formation than the ‘No-host control’ (n=48-54, Kruskal Wallis rank sum test, df=2, p=0.004), Dunn test comparison No-host control vs Pathogen only p=0.003). **E)** Overall, there was a significant positive correlation (n=16-18, Spearman’s rank correlation rho, p=0.005) between biofilm formation and virulence phenotype of the evoved pathogens. When broken down by pathogen evolutionary history, the ‘Pathogen only’ group showed a significant positive correlation (n=17, Spearman’s rank correlation rho, p=0.045) between these two phenotypes.

After 15 passages, we assayed each evolved pathogen for virulence (host killing) and colonisation ability. For all evolved pathogens, host mortality due to infection was first measured in the absence of the ancestral microbiota. The pathogen that evolved in isolation caused significantly higher mortality than the no-host control (fig 3A). In contrast, the pathogen that had evolved with ancestral microbiota caused similar levels of mortality to the no-host control (fig 3A). No significant difference in virulence was found between the pathogen that evolved in isolation compared to the pathogen that evolved with ancestral microbiota. The observed increase in virulence of the pathogen evolved in isolation compared to the no-host (ancestral) control was not mediated by increased colonisation, since all evolved pathogens exhibited a similar ability to colonise nematode hosts (fig 3B).

### *agr* activity and biofilm formation contribute to the increase in evolved *S. aureus* virulence

We found that *in vivo* RNAIII expression was on average 27-fold higher in the pathogen evolved alone compared to the no-host control (fig 3C). *agr* is a well-characterised global regulator of staphylococcal virulence ^43–45^, and RNAIII is the effector of target gene regulation ^46^. Activation of *agr* results in increased expression of secreted proteins involved in virulence ^47^. We simultaneously tested RNAIII expression in the ancestral pathogen in the presence and absence of ancestral microbiota, to see if upregulation of *agr* explained the original increase in virulence shown in fig 1A. However, no difference was observed in *agr* expression in the pathogen that infected microbiota-colonised hosts (fig S1a).

We also assessed the ability to form *in vitro* biofilm of each of the evolved pathogens. Biofilm formation is a well-studied aspect of virulence in *S. aureus* ^48–50^. In *C. elegans*, the formation of staphylococcal biofilm contributes to the establishment of infection via attachment to host cells, and to resistance to host immune factors ^51^. Thus, biofilm is an important contributor to virulence in this nematode model. This phenotype was measured using a 96-well plate assay, in which biofilms were stained with crystal violet after 24 hrs bacterial growth and quantified by optical density (595nm). In comparison to the no-host control, the pathogen that evolved in isolation exhibited significantly higher biofilm formation (fig 3D), which positively correlated with its killing ability (fig 3E). Further research, involving knock-out bacterial mutants, is needed to demonstrate a causative relationship.

### No difference in virulence of evolved pathogens in hosts colonised by ancestral microbiota

When hosts were colonised by the ancestral microbiota, all evolved pathogens caused comparable levels of host mortality (fig S1b). All evolved pathogens also became better at competing with host microbiota. Evolved pathogen load was consistently higher in the presence compared to the absence of host microbiota (fig S1c), and microbiota load was significantly reduced in infected compared to uninfected hosts (fig S1d). This result suggests that adaptation to the lab environment (not just host environment) favoured a competitive fitness advantage for *S. aureus*.

### The effect of evolved pathogens on microbiota community assembly is host-mediated

We investigated how the evolved pathogens’ competitive colonisation advantage affected microbiota community structure. We conducted 16S sequencing on nematodes colonised by ancestral microbiota and each evolved pathogen. We found that alpha diversity and composition of the ancestral microbiota community did not differ across each of the evolved pathogens, within *in vitro* or *in vivo* samples (figs S2-3). Composition and alpha diversity differed significantly between paired *in vitro* and *in vivo* samples however (fig S2), indicating a strong host-mediated effect of infection on microbiota community assembly, as observed in the ancestral state. *In vivo*, MYb10 dominated across all groups, followed by MYb71 and MYb11. BIGb393 and BIGb0170 were mostly excluded from the community, while CEent1 and JUb66 were consistently present at low abundances (fig S3a). *In vitro*, CEent1 dominated, making up 75% or more of the community in each replicate. BIGb393 and MYb71 were mostly excluded, while MYb11 was present at low abundance in around two thirds of replicates. MYb10 was present in most replicates at low abundance. JUb66 and BIGb0170 were consistently present across replicates but also at low abundances (fig S3b).

### Genetic diversification of *S. aureus* is slower in the absence of microbiota

We next evaluated the molecular basis for differences in evolved pathogen virulence using whole genome sequencing. For all six replicates of each evolved pathogen, forty clones were pooled for population sequencing. Simultaneously, two clones were individually sequenced per replicate. Populations and clones were sequenced for passages 10 and 15 of the evolution experiment, to allow characterisation of evolutionary changes over time.

Variant calling was conducted from sequencing data to identify signatures of genomic evolution. We filtered out ancestral and no-host control SNPs from evolved lineages, to focus our analysis on *de novo* mutations.

Whole genome sequencing of clones at passage 15 revealed that significantly fewer mutations, specifically non-synonymous mutations and indels (fig. S5a) occurred in pathogen clones evolved in isolation (fig 4A). Any SNPs present at >=25% frequency within pooled populations of each replicate, after filtering of ancestral and no-host control SNPs, were classified as targets of selection. Table S1 shows the full list of genes under selection across evolved pathogen replicates at passage 15. We used the genome comparison tool ACT ^52^, BLAST searches for homologous proteins and literature searches to ascertain the function of each gene under selection. Genes were categorised by broad functional category (metabolism, virulence or adherence) based on their primary function. Genes under selection in the pathogen that evolved alone with possible links to staphylococcal virulence include *purR*, *icaC*, *gltD*, *clfA*, and *dltB*, along with an intergenic region close to the *mnh* operon. However, no clear pattern emerged to indicate a genomic basis for the change in virulence of pathogen populations in this group, with different genes under selection in each replicate. These results suggest that virulence under selection in our experiment is based on multiple genes, as found in other pathogens with broad host ranges ^53–55^. A number of SNPs were located in genes with links to biofilm formation. Where the pathogen evolved with ancestral microbiota, 4 SNPs across 3 replicates had links to biofilm formation, along with 4 SNPs across 3 replicates in the pathogen that evolved alone (see table S1). In the latter, these genes were *icaC*, *gltD*, *clfA* and an intergenic mutation in the ribosome binding site upstream of the gene SAS_RS08335, encoding *apt*. Molecular evolution in pathogens that evolved with ancestral microbiota initially led to rapid diversification. We calculated genetic distance among replicates for each of the *in vivo* evolved pathogens using population-level sequences. At passage 10, replicate populations in which the pathogen evolved with ancestral microbiota were more genetically distant from each other compared to replicates of the pathogen that evolved alone (fig 4B). However, by the final passage, the genetic distances among replicates were comparable between the two evolved pathogen groups (fig 4B). This finding suggests that without the additional selective pressure of the microbiota, pathogen genomic diversification is slower.

**Figure 4.**
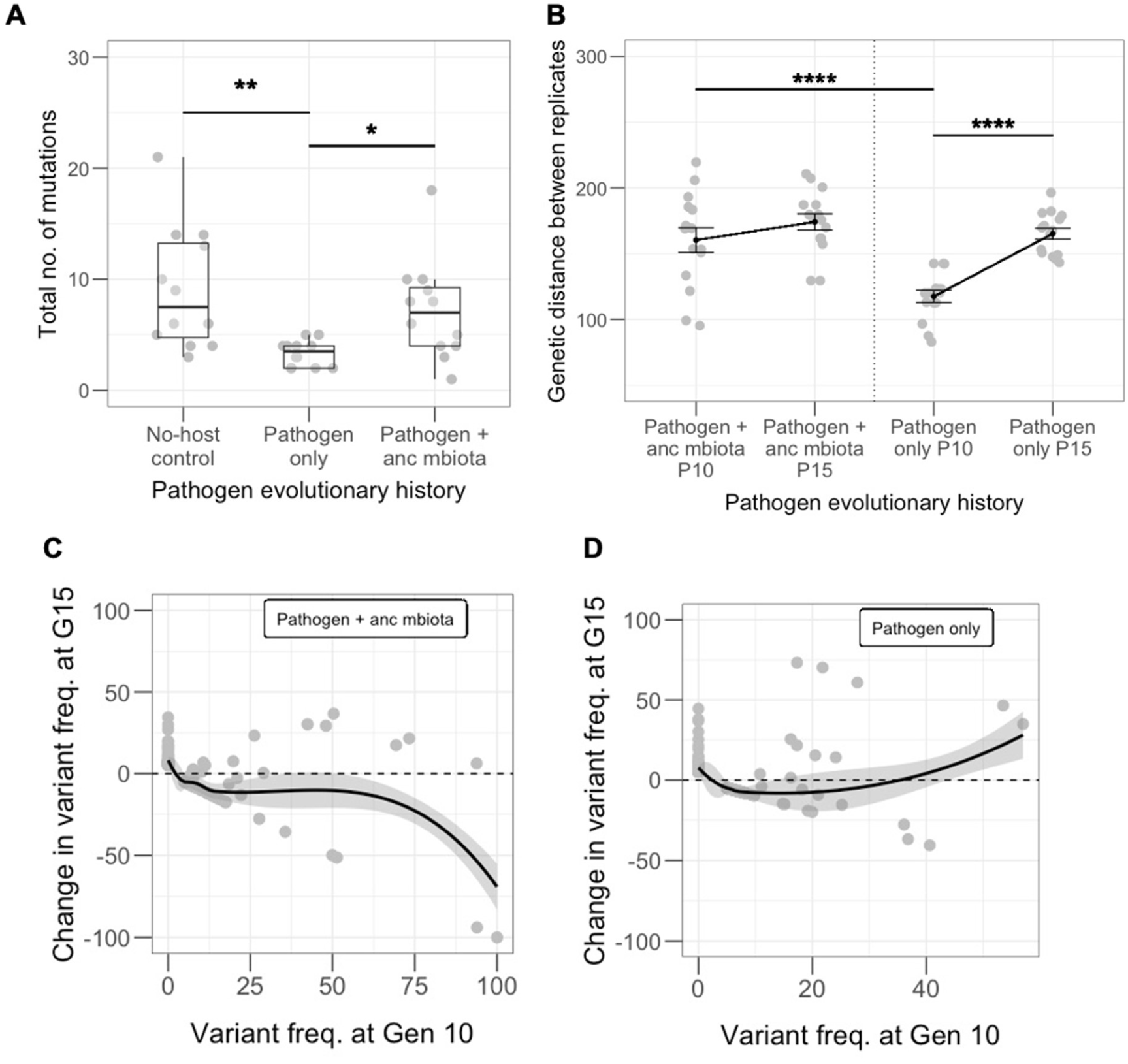
**A)** After 15 passages, significantly fewer mutations occurred in clones of the ‘Pathogen only’ group compared to clones that evolved alongside the microbiota (n=12, Kruskal Wallis rank sum test, df=2, p=0.002, Dunn test comparisons: No-host control vs Pathogen only p=0.003, Pathogen only vs Pathogen + anc mbiota p=0.017). **B)** Genetic distance between replicate populations within each evolved group at generations 10 and 15. Significantly less genetic distance was found between replicates of the ‘Pathogen only’ group at generation 10 compared to generation 15 (n=15, ANOVA, f=15.28, df=3, p<0.0001, Tukey HSD comparison between Pathogen only P10 vs P15 p<0.0001) and compared to generation 10 of the ‘Pathogen + anc mbiota’ group (TukeyHSD p=0.0001), indicating a slower rate of diversification in populations of the ‘Pathogen only’ group. **C)** Negative frequency-dependent selection was observed when the pathogen evolved alongside the ancestral microbiota (n=205, linear regression, f=66.9, df=1, p<0.0001), but **D)** not in the ‘Pathogen only’ group (n=301, linear regression, f=3.16, df=1, p=0.076).

We calculated the genetic distance of replicates in each group from the ancestor. No differences were observed in distance from the ancestor, indicating no difference in the overall rate of evolution between the two groups (fig S5b). We also analysed nucleotide diversity across passages 10 and 15 between the two groups and found no significant difference (fig S5c). The frequency of mutations ranged from 5-100%, of which there were generally more nonsynonymous than synonymous SNPs across treatments at each time point (figs S5d-f). Across populations, there was variation in terms of the frequency of nonsynonymous vs. synonymous SNPs. For example, most mutations in pathogen only P15 population six were at <25% frequency, whereas populations one and three of the same treatment and time point had many mutations at >50% frequency (figs S5g-h). Across the genome, there were more nonsynonymous than synonymous SNPs (fig S5i). One region in the genome exhibited a high number of mutations at moderate frequencies, but because this occurred across all treatments at both time points, these mutations likely arose from general passaging and not any particular treatment. Table S2 breaks down the number of SNPs in genes and intergenic regions for each population within each treatment across the two time points. On average, each treatment had similar densities of genic mutations by passage 15.

### Negative frequency-dependent selection acts on *S. aureus* in hosts with a microbiota

We examined the modes of selection operating throughout the experiment, to further characterise pathogen evolutionary dynamics. For the pathogen that evolved with ancestral microbiota, approximately 30% of genes under selection fluctuated in frequency throughout the experiment (fig S5j). In contrast, only 12.5% of genes under selection fluctuated in the pathogen that evolved alone (fig S5k). Consequently, we looked for signatures of negative frequency-dependent selection using a standard approach^56,57^, in the pathogen that evolved in hosts with ancestral microbiota. Negative frequency-dependent selection (NFDS) occurs when rare variants are selected for within a population, thus individuals with the most common genotype are at a selective disadvantage^58,59^. In a direct test of NFDS on pathogens evolved with ancestral microbiota, we found a significant negative linear relationship between a mutation’s change in frequency across the experiment and its frequency at the mid-point (fig 4C), with 58% of alleles increasing in frequency from passage 10 to 15. Pathogens evolved alone were not under NFDS (fig 4D). This finding suggests that when in competition with the microbiota, rare pathogen genotypes are at a selective advantage.

### The microbiota passaged in isolation facilitates pathogen virulence to a greater extent

We hypothesised that microbiota communities maintained within host populations over evolutionary time would dampen virulence of the ancestral pathogen. After passaging the microbiota through 15 host generations (fig 2B), we found that microbiota passaged in infected hosts did not diminish virulence of the ancestral pathogen (fig 5A). However, microbiota passaged through uninfected hosts drove significantly more host killing by ancestral *S. aureus* compared to the no-host control (fig 5A).

Microbiota load in the face of ancestral pathogen infection was comparable across all evolved groups (fig 5B – circular symbols) as was pathogen load (fig 5B – triangular symbols). In all cases, the ancestral pathogen was more competitive than the microbiota, colonising nematodes to a significantly greater extent than the evolved microbiota communities (fig 5B). *agr* expression in the ancestral pathogen was measured in the presence of each evolved microbiota community by qRT-PCR, but no significant difference was observed (fig S6). Therefore, the higher level of host killing from *S. aureus* in the presence of microbiota that evolved alone is not due to higher expression of virulence factors by the pathogen. Microbiota passaged in uninfected hosts caused 2% average mortality in the absence of the pathogen. However, this was not significantly different from the no-host control (fig S7). Thus, the observed increase in host mortality in the presence of *S. aureus* could not be explained by direct killing from the microbiota community maintained across generations.

### Passage in infected hosts significantly reduces microbiota community stability

To examine how *S. aureus* affected the community composition of passaged microbiota populations, we conducted metagenome sequencing on microbiota communities from passage 15. We found that species richness was significantly higher in the no-host control compared to all other communities (fig 5C). However, species evenness in the no-host control was significantly lower (fig 5C). Composition of the microbiota communities differed significantly between replicates of each evolved group (fig 5D). MYb10 was enriched in the microbiota that evolved in infected hosts, while MYb71 and BIGb0170 were enriched in the no-host control. In the presence of the pathogen, the microbiota became significantly more dispersed compared to all other microbiota communities (fig 5E). Specifically, pathogen infection drove the microbiota to form two distinct profiles, dominated either by MYb10 and JUb66, or MYb10 and CEent1 (Figs. 5D & 6A).

**Figure 5.**
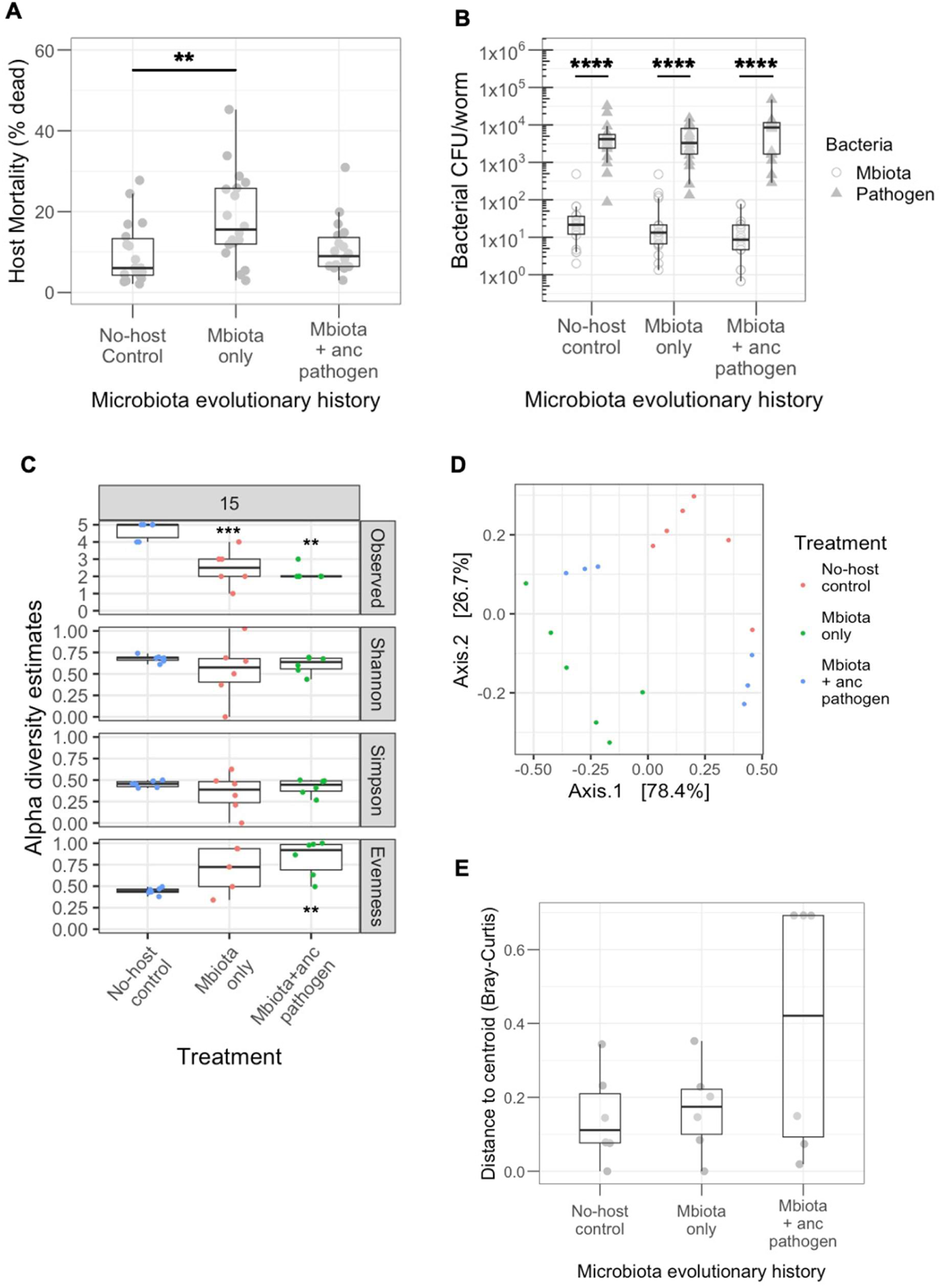
All data shown represents the 15th (final) evolved passage. **A)** The microbiota that evolved alone facilitated the virulence of ancestral *S. aureus* to a significantly greater extent than the no-host control evolved microbiota (n=18, Kruskal Wallis rank sum test, df=2, p=0.01, Dunn test comparison No-host control vs Mbiota only p=0.01). **B)** Colonisation of *C. elegans* was comparable across all three evolved microbiota lineages, which all colonised the worms to a significantly lesser extent than the ancestral pathogen (n=35-36, linear regression, f=87.73, df=5, p<0.001, post-hoc comparisons using ‘emmeans’ package: Mbiota CFU/worm vs Pathogen CFU/worm p<0.0001 in each evolved group). **C)** Alpha diversity estimates of each evolved microbiota community. Stars indicate significant differences from the no-host control in each graph. ‘Mbiota only’ has significantly reduced species richness (n=12, linear regression, f=20.61, df=1, p=0.001), as does ‘Mbiota + anc pathogen’ (n=12, linear regression, f=55.38, df=1, p=0.003). ‘Mbiota + anc pathogen’ also has significantly reduced evenness (n=12, linear regression, f=30.86, df=1, p=0.0002) **D)** Principal component analysis showing beta-diversity of evolved microbiota communities. Community composition differed significantly between replicates of each evolved group (PERMANOVA Bray Curtis, p=0.0001). In particular, the microbiota that evolved with ancestral pathogen is significantly more dispersed compared to the microbiota that evolved alone (PERMANOVA Bray-Curtis, f=21.26, df=1, p=0.001). **E)** Where the microbiota evolved with the ancestral pathogen, evolved communities are significantly more unstable compared to the ‘Mbiota only’ group (PERMANOVA, f=21.158, df=1, p=0.001).

Pathogen infection altered competitive dynamics within the microbiota. Having been passaged in hosts infected by the ancestral pathogen, strains CEent1 and JUb66 were never found to co-exist at passage 15. In contrast, MYb10 was always maintained in these replicates (fig 6A). The opposite dynamic was seen in replicates of the microbiota evolved in uninfected hosts (fig 6A), whereby CEent1 and JUb66 regularly co-existed, but MYb10 was mostly absent from the community. We found that CEent1 and JUb66 were phylogenetically closely related (fig 6B) and had a high degree of metabolic resource overlap (MRO) (fig 6C). A high degree of resource overlap, as found between CEent1 and JUb66, indicates strong competitive interactions for nutritional resources. Manipulation of the *in vivo* nutritional environment is needed to further characterise these interactions.

**Figure 6.**
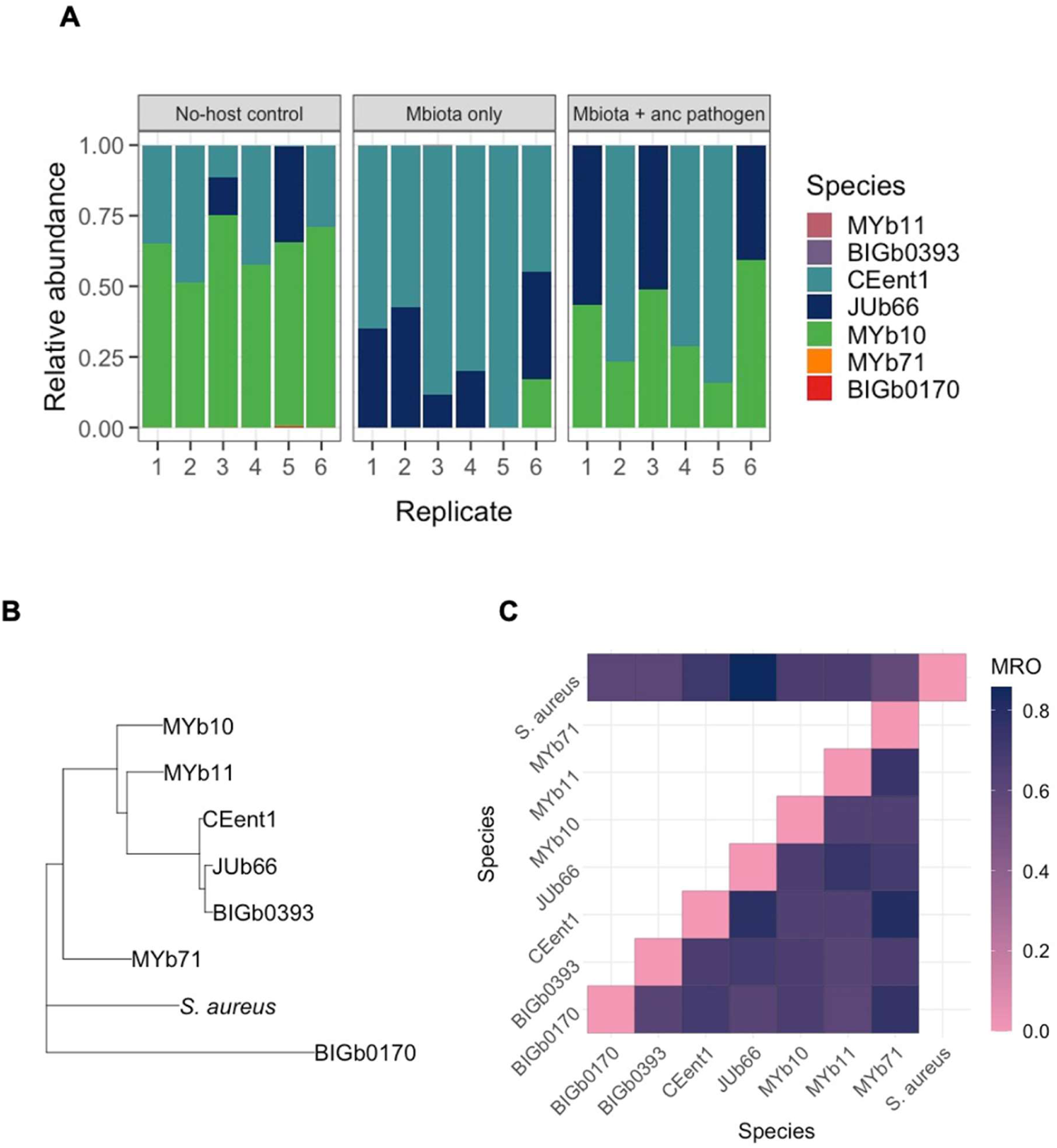
**A)** Community composition of each replicate population within the three evolved microbiota groups at passage 15. MYb10 is significantly enriched in the microbiota that evolved in infected hosts (Wilcoxon rank sum test, p=0.0002). MYb71 and BIGb0170 are enriched in the no-host control (p=0.003 and 0.001, respectively). **B)** Phylogram depicting relatedness of the seven microbiota species and *S. aureus*. **C)** MRO between bacterial strains and pathogen pairs. JUb66 and CEent1 are phylogenetically close and have higher MRO, indicating higher competition for resources between these two species.

### Host microbiota has strong cross-feeding interactions with *S. aureus*

We investigated cross-feeding interactions between microbiota components to identify which resources were the source of competition. BIGb0170 and BIGb0393, both as metabolic donors to *S. aureus* of Fe2+ and O2, have high smetana scores of 1 (a smetana score of 1 indicates absolute certainty on the cross-feeding interaction). CEent1 can serve as a donor of copper and Fe2+ to *S. aureus* (smetana score=1). MYb10 and MYb71 can donate phosphate and copper to *S. aureus*, respectively (smetana scores=1). Conversely, *S. aureus* can donate O2 to MYb10 and MYb71 (smetana scores=1), as well as donating phosphate to MYb71 (smetana score=1) and L-Cysteine to MYb10 (smetana score=0.9). Within the microbiota, cross-feeding interactions are much weaker, shown by smaller smetana scores for microbe pairs (see supplementary file 1). Particularly, we found that JUb66 and CEent1 are exclusively donors and do not receive metabolites from other microbiota members, which might exacerbate their competition for nutrients in *S. aureus*-infected hosts.

### Short-term pathogen evolution

We tested whether transplanting a host-adapted microbiota community could alter the trajectory of within-host pathogen evolution over a short timescale. We introduced microbiota evolved in uninfected hosts (from passage 10) into the group in which pathogens were evolving alone. Both microbiota and pathogen in this newly created group, likewise consisting of six replicates, evolved for the remaining five passages alongside the original evolving groups (fig S8). Comparing the virulence of *S. aureus* at passage 15 between this group and the original pathogen that evolved alone, we did not find any significant differences in host mortality from infection (fig S9). Introduction of microbiota adapted to uninfected hosts therefore did not significantly alter the trajectory of *S. aureus* virulence on this time scale.

## Discussion

Most microbes face constant warfare with others ^60,61^, which may be heightened during infection by a pathogen. Competition between pathogens ^62^, or within microbial symbiont communities ^60^, can be a major selective force shaping virulence. Here, upon initial invasion of nematode hosts, *S. aureus* displaced resident microbiota (in terms of number and diversity) worsening disease outcome, comparable to results observed with other bacterial pathogens ^63^. We subsequently found the evolutionary paths of pathogens and host microbiota varied depending upon their competitive opportunities within nematode hosts.

In our experiment, competition with host microbiota maintained pathogen virulence over evolutionary time at levels seen in ancestral controls. The dominance and high virulence of *S. aureus* in microbiota-colonised hosts could be likened to ‘super-infections’ whereby a virulent pathogen can take over a host already infected by a less virulent one. Nowak and May ^18^ found that superinfections can maintain highly virulent pathogen strains across evolutionary time if virulence is equated with within-host competitive advantage. The high pathogen load and high level of host harm caused by infection in nematodes with natural microbiota may thus limit any movement in optimal virulence. However, we found competition with host microbiota caused a burst of pathogen molecular evolution. Superinfection can lead to complex evolutionary dynamics involving heteroclinic (oscillating) cycles of genetic mutations within a population ^18^, which can be driven by NFDS generated from species interactions ^64,65^. We found that pathogens evolving amidst nematode microbiota were more likely to experience NFDS at the genomic level. *S. aureus* is a relatively clonal organism, as such point mutations are the more common source of new alleles in this species^66^. Greater genetic divergence was also observed among replicate evolved populations in this group, with pathogen populations evolving in isolation grouped together genotypically. NFDS has been found to similarly cause population diversification in other microbial species interactions ^67,68^.

Pathogen evolution in isolation favoured a two-fold increase in host mortality from infection compared to the ancestral controls. This outcome is common in serial passage experiments, in which transmission is guaranteed by experimental design, therefore reducing the cost of high virulence ^69^. The increase in virulence in the pathogen that evolved alone could be due to selection on a combination of virulence factors. Namely, we found that biofilm production and the expression of *agr,* a global virulence regulator, were higher in the pathogen that evolved alone compared to the no-host control. The ability to form a biofilm can improve a pathogen’s ability to colonise host tissues and resist immune clearance ^51,70^ This trait has been a target of selection during evolution experiments and is an important aspect of pathogen virulence in *C. elegans* ^71^*. agr* has been shown to contribute to staphylococcal infection in a range of host models ^40,44,72^, with *agr*-deficient mutants exhibiting reduced virulence ^44,73,74^. Selection on *agr* has been revealed to impact virulence evolution in *S. aureus* ^75^. It controls expression of a range of secreted toxins, including the alpha, beta and delta haemolysins, toxic shock syndrome toxin and staphylokinase ^43^. High *agr* activity is often associated with reduced biofilm formation – or dispersion and detachment of staphylococcal biofilms ^70,76,77^. The fact that quorum-sensing activates *agr* at high cell densities suggests its activity may vary in our system as infection progresses ^78^. Perhaps a biofilm forms in the host gut initially, with *agr* expression induced as pathogen densities increase, allowing cells to detach from biofilms and disperse within the site of infection ^78^.

Pathogen presence during *in vivo* microbiota passages strongly shifted community dynamics. Infection is often indirectly associated with dysbiosis of the microbiota ^9,79,80^. Here, we found a direct link between *S. aureus* infection and increased nutrient competition between resident microbes, with destabilising effects on community composition over evolutionary time. Whilst *Acinetobacter gouilliae* (MYb10) can be a recipient of nutrients during cross-feeding with *S. aureus,* and thus maintained in every replicate, pathogen infection prevented the coexistence of two closely related microbiome species. We observed the stochastic competitive exclusion of either *Enterobacter hormaechei* (CEent1) or *Lelliottia amnigena* (JUb66) within all replicates of microbiota communities passaged with the pathogen. Similar dynamics have been observed with *Clostridioides difficile*, whereby microbiota community structure is altered during infection due to changes induced in the nutritional landscape ^81^. Our findings fit with the Anna Karenina Principle, whereby dysbiotic microbiota communities differ more in community composition compared to healthy (eubiotic) communities ^82^. We find that destabilization arises in infected hosts across both ecological and evolutionary time. This pattern suggests that infected hosts are unable to regulate their microbiota community ^82–84^, with microbiota composition lacking a steady state in the face of pathogen invasion ^85,86^.

We found that higher disease severity was caused when worms were colonised by microbiota passaged in isolation and infected by ancestral pathogens. A parallel can be drawn between our findings and the high levels of virulence often observed when a pathogen jumps into a new host. *S. aureus* in particular has a wide host range ^32–34^ and is known to jump between animal species ^87–90^. Upon invading a new host, *S. aureus* will also encounter a new microbiota with which it does not share an evolutionary history.

Competition with the resident microbiota community might therefore contribute to more severe disease outcomes from novel *S. aureus* infection. Having previously travelled on separate evolutionary paths, we found invading (ancestral) *S. aureus* and host-adapted resident microbiota exhibit greater antagonism upon first meeting. Considerable research has been conducted to characterise interactions between *S. aureus* and host microbiota communities ^91–94^. Future investigations seeking to understand disease severity from *S. aureus* infections, and other generalist pathogens with spillover potential ^95,96^, should consider the role of competition with host microbiota during these novel interactions.

Patterns of infectious disease distribution and transmission are predicted to undergo considerable changes in coming years ^97^. Climate change, biodiversity loss, and associated human activities are providing more opportunities for pathogens to jump between animal species and humans ^98^. Our study reveals that when pathogens infect new hosts, competitive dominance over microbiota sparks pathogen molecular diversification, while virulence evolution is limited. Similar pathogen genome dynamics have been uncovered during coevolution whereby changes in host genetic background, but not microbiome communities, generates fluctuating selection ^99^. Moreover, the ecological factors imposing selection on microbiomes are increasingly being uncovered ^100,101^. For host microbiomes herein, infection by a novel pathogen was a perpetually destabilising force. Such variability and stochasticity in microbiome composition highlights the difficulty of predicting infection outcome during epidemics of new pathogens. Ultimately, understanding the mechanisms of host microbiota-pathogen interactions may help explain infection outcomes now and across evolutionary time both in wildlife and humans.

## Methods

### Species and strains used

The *C. elegans* homogenous N2 line was used as a non-evolving host throughout the experiment. *C. elegans* is an established model host for microbial colonisation and pathogenesis ^40,102,103^. It is a natural predator of bacteria ^104–106^, but it can also be colonised by various pathogen and commensal species ^40,103,107^. Recent work has characterised the resident microbiota of the *C. elegans* gut, resulting in formation of the CeMBio collection of microbes, which represent the most common components of its natural gut microbiota ^42^. As a non-evolving host in this experiment, worm populations were freshly resurrected at each passage to prevent the accumulation of *de novo* mutations.

The ancestral microbiota community originated from the CeMBio collection and consisted of seven bacterial species, all of which are representative of a naturally associated *C. elegans* microbiome ^42^. The seven species were: *Pantoea nemavictus* (BIGb393), *Lelliottia amnigena* (JUb66), *Sphingobacterium multivorum* (BIGb0170), *Enterobacter hormaechei* (CEent1), *Acinetobacter guillouiae* (MYb10), *Pseudomonas lurida* (MYb11) and *Ochrobactrum pecoris* (MYb71). Each component was selected based on their ability to grow on XLD (Xylose Lysine Deoxycholate) media to distinguish them from *Staphylococcus aureus*. *Escherichia coli* strain OP50 was used as a food source for worms which were not fed microbiota.

Strain MSSA476 (GenBank: BX571857.1) was used as the pathogen, sourced from the University of Liverpool. Previous work from our lab and others has shown that *S. aureus* is an effective pathogen of *C. elegans*^7,25,40^. Accumulation of *S. aureus* in the nematode intestine has been correlated with increased host mortality^40^, causing enterocyte effacement, intestinal epithelium destruction, and degradation of internal organs ^39^. Expression of key virulence determinants such as the quorum-sensing global virulence regulatory system *agr*, the global virulence regulator *sarA*, the alternative sigma factor σB, alpha-hemolysin, and V8 serine protease have all been implicated as requirements for full pathogenicity in this nematode model ^41,108^. MSSA476 is an invasive community-acquired methicillin-susceptible isolate originally isolated from a human ^109^.

### Worm and bacterial growth conditions

Unless specified otherwise, all microbiota strains and OP50 were cultured in 10ml LB broth, inoculated with a single colony, and incubated at 25°C (shaking 150 rpm). *Staphylococcus aureus* was cultured in Todd Hewitt broth (THB), inoculated with a single colony, and incubated at 30°C (shaking 150 rpm). On agar plates, microbiota and OP50 were cultured at 25°C on LB agar to obtain single colonies, whilst *S. aureus* was cultured at 30°C on Tryptic Soy Agar (TSA). Evolved bacterial populations used in follow-up host mortality assays were inoculated into 10ml liquid media directly from the freezer stock. All other incubation conditions remained the same.

To isolate and sterilise eggs, gravid *C. elegans* worms were suspended in 6ml M9 buffer containing 0.1% Triton-X (M9-Tx) and treated with 1ml bleach (1:1 mix of 5M sodium hydroxide and sodium hypochlorite). Worms were incubated in bleach at room temperature for up to 10 minutes, centrifuged (2 mins, 400 x g) and washed twice with M9-Tx. Worms were then resuspended in plain M9 and incubated at 20°C overnight (shaking 150rpm). This allowed eggs to hatch and synchronised the population, such that all worms were arrested at the L1 larval stage. Hatched worms were transferred onto NGM plates seeded with 600µl OP50 liquid culture and incubated at 20°C for 48 hrs. L4 worms were then transferred to infection plates as described below. To maintain worms throughout the evolution experiment, a chunk of NGM with worms was cut out with a sterilized spatula and transferred to a fresh NGM plate seeded with OP50, then incubated at 20°C.

### Evolution experiment design

The evolution experiment consisted of six different groups of evolving bacteria; four *in vivo* treatments and two no-host controls, each replicated six times (see fig 2B). Briefly, both pathogen and microbiota were evolved alongside their ancestral opponent (‘Pathogen + anc mbiota and Mbiota + anc pathogen) and in isolation (‘Pathogen only’ and ‘Mbiota only’). No-host controls consisted of the pathogen and microbiota each evolving *in vitro* in isolation. *In vivo* evolving lines were passaged 15 times through non-evolving *C. elegans* populations.

No-host controls were passaged alongside, acting as proxies for the ancestor(s) and controlling for lab adaptation.

Ancestral bacteria were re-introduced at each passage where relevant. For groups including a microbiota, the *C. elegans* food source was the 7-species microbiota community; each species was cultured individually, then standardised to the optical density (OD600) of the species with the lowest growth and pooled into one tube. For the ‘Pathogen only’ group the food source was *E. coli* OP50, also standardised in line with the microbiota, to ensure no disparity in the amounts of bacteria available as food. 600µl of food (pooled microbiota or OP50) was plated onto 9cm NGM plates. Approximately 1000 hatched L1 worms were added to each plate, except the ‘No-host control’ group, and all plates were incubated at 20°C for 48hrs. A further 100µl of the pooled microbiota community was spread onto additional NGM plates for use as substitute infection plates for the ‘Mbiota only’ group. This ensured worms were fed the same microbiota community throughout the cycle (these plates were incubated for 24hrs at 20°C then at 4°C until used at the infection step). To create infection plates for groups including the pathogen, *S. aureus* cultures were made and, following incubation, 100µl was spread onto 9cm TSA plates. These plates were incubated for 24hrs at 30°C.

After 48hrs incubation on food, worms were washed off plates with 850µl M9-Tx. Worms were then transferred to cut-off 1000μl filter tips within 1.5ml tubes and washed twice with 250µl M9-Tx (centrifugations were 2 mins, 290 x g). They were resuspended in 100µl M9-Tx and transferred to either *S. aureus* infection plates or to pre-prepared microbiota plates.

Infection and microbiota plates, along with the untreated no-host controls, were incubated at 25°C for 24hrs.

A gentle bleaching protocol was conducted as described in ^42^) to remove any surface-adherent bacteria from the worm cuticle. This was to ensure that, as far as possible, only gut bacteria were extracted from the host for passage. Worms were transferred in 100 µl M9 buffer (containing 0.025% Tx) and crushed using a Bead Bug Microtube Homogeniser (Benchmark Scientific) for 3 mins at 320 rpm. Approximately 100 worms remained in each replicate, representing 10% of the original population.

Ten-fold serial dilutions were made of the crushed worm solution. 10^-2^ dilutions were plated onto XLD selective media to isolate the microbiota, while 10^-4^ dilutions were plated onto MSA to isolate the pathogen. For the no-host controls, a 1µl inoculation loop was used to transfer a sample from the bacterial lawn onto XLD for microbiota and MSA for pathogen. XLD plates were incubated for 48hrs at 25°C, while MSA plates were incubated at 30°C for 24hrs.

To passage bacteria to the next host generation, 100 colonies were picked from selective plates for microbiota and pathogen, for inoculation into 10ml LB or THB, respectively. For the no-host controls, an inoculation loop was used to take a sample from the bacterial lawn, which was likewise inoculated into 10ml liquid media. Aliquots of these liquid cultures, were archived in 25% glycerol at -80°C. The liquid cultures were then used to make the next passage’s food/infection plates as described above. OP50 and evolving microbiota cultures were standardised to the ancestral species with the lowest growth from passage 2 onwards, to ensure the same amount of food was plated for all lineages.

### Host mortality assays

To evaluate pathogen virulence in the presence and absence of microbiota, the level of host mortality was assessed upon exposure to *S. aureus*. Infection/microbiota plates were made as described above. However, 60µl *S. aureus* was plated onto smaller 6cm TSA plates, and for the evolved microbiota only 400µl liquid culture was plated onto NGM. After 24hrs incubation of infection plates, they were randomly divided in half and the number of live and dead worms in one half were counted. Raw data was multiplied by 2 to account for the whole plate. The proportion of dead worms was calculated as a measure of virulence. Three to six technical replicates were conducted for each sample.

### Bacterial growth assays

To assay bacterial colonisation of the *C. elegans* gut, the evolution experiment protocol was followed up to the ‘soft bleach’ step. After the ‘soft bleach’ step, five worms were sampled from each replicate and transferred to 100µl M9. This solution was crushed as per the protocol described above and serial dilutions prepared in M9. For each sample at each dilution, three 10µl spots were plated onto MSA/XLD agar. Averages of three 10µl spots were calculated using the optimal dilution, from which CFU/worm was calculated.

Growth of the microbiota and pathogen, both individually and in competition, was assayed *in vitro*. Each species was cultured separately overnight, standardised to the same optical density (OD600) and pooled as described above. 1.5µl of pooled microbiota and/or 1.5µl pathogen was added to 1.5ml fresh LB broth. These cultures were incubated for 24 hrs at 25°C to replicate the infection conditions of the *in vivo* experiments. Cultures were then serially diluted 10-fold and spots plated as described above.

### Pathogen biofilm assay

Biofilm formation of ancestral and evolved *S. aureus* was assessed. Evolved pathogens from passage 15 were inoculated directly from freezer stocks into 10ml THB broth and incubated at 30°C with shaking for 24 hrs. Cultures were then diluted 1:40 into 100µl THB containing 0.5% glucose in a 96-well plate. Wells at the edges of the plate were filled with sterile water to prevent evaporation of the samples, and the plate was incubated statically for 24 hrs at 30°C. To further reduce the risk of evaporation, plates were incubated inside a plastic box lined with wet paper towel. Positive (*S. aureus* MSSA476) and negative (*S. aureus* strain LAC) controls were included in this assay. After incubation, samples were washed once with sterile water, stained with 150µl 0.5% crystal violet for 30 mins at room temperature, washed once more with sterile water and finally resuspended in 200µl 7% acetic acid. OD(595) readings were taken for each sample using a plate reader.

### 16S and metagenomic sequencing of microbiota

For 16S sequencing, bacterial DNA was extracted from crushed worms using a ZymoBIOMICS DNA Miniprep kit (Zymo) according to the manufacturer’s protocol. The V3- V4 regions of bacterial 16S rRNA were amplified using the universal primer pair 341F (5’- CCTACGGGNGGCWGCAG-3’) and 805R primer (5’-GACTACHVGGGTATCTAATCC-3’).

PCR amplicons were sequenced on the Illumina Miseq platform using 2 x 300bp v3 chemistry by the Integrated Microbiome Resource at Dalhousie University, Canada. FastQC ^110^ and MultiQC ^111^ were used for initial visualization of read quality, primers were removed using Cutadapt ^112^. Paired-end reads were joined using vsearch ^113^. All low-quality reads were then filtered using default quality thresholds before starting the Deblur ^114^ workflow to denoise and classify sequences into amplicon sequence variants (ASVs). Trimming length was determined as 400 bp after manually viewing the quality plot. As full-length 16S rRNA sequences for the seven microbiota species were well- characterized ^42^, the obtained sequencing reads were processed through the closed-reference OTU picking pipeline in QIIME2 ^115^. To build the reference, full-length 16S sequences were downloaded for the seven microbiota species and converted to a qza-formatted reference file for processing by QIIME2. Taxonomy of the resolved ASVs was assigned by clustering ASVs to the customized reference with 99% similarity thresholds.

The relative abundance table was rarefied to the minimum sample size and alpha-diversity indices – Richness, Shannon diversity, Simpson index and Evenness were computed using the R package phyloseq. For beta diversity, Bray-Curtis dissimilarity was calculated based on rarefied relative abundance, using the R package phyloseq. Permutational analysis of variance (PERMANOVA) was conducted with 9999 replications on each distance metric to evaluate differences in the microbiome structure and composition between treatments using the R package vegan ^116^. Microbiome dispersion was calculated using the betadisper function in R. Differences in microbiome dispersion between different treatments were tested using a permutation test.

16S rRNA sequences were downloaded for each of the seven microbiota species and the pathogen. Multiple sequence alignment (MSA) was performed using MUSCLE ^117^ and the MSA was used to reconstruct a maximum-likelihood tree in IQTREE (v1.6.11) ^118^. Tree reconstruction was performed with 1000 ultrafast bootstrap replicates and the SH-like approximate likelihood ratio test (“-bb 1000 -alrt 1000”). The best-fit model “TN+F+G4” was selected by ModelFinder ^119^ based on the Bayesian information criterion (BIC). The phylogenetic distance matrix was generated from the consensus tree using the ‘cophenetic’ function in R.

Metagenome sequencing of microbiota was conducted on passage 15 of the evolved microbiota communities. 10ml LB cultures were inoculated straight from the evolved microbiota freezer stocks, by taking a sample of frozen bacteria with a 1µl loop. Cultures were incubated overnight at 25°C, with shaking. Cultures were then pelleted by centrifugation, resuspended in 1ml PBS and incubated with 25 units mutanolysin overnight at 37°C, with shaking. Samples were then digested with 0.5µg RNase A for 15 minutes at room temperature. DNA was extracted using the High Pure PCR Template Preparation Kit (Roche) as per the manufacturers protocol.

Sequencing was conducted by the Centre for Genomic Research at the University of Liverpool, UK. Extracted DNA was prepared for Illumina short-read sequencing using the NEBNext Ultra II Kit, using 1/2 volume reactions. DNA was sequenced by Illumina NovaSeq using S4 chemistry (Paired-end, 2x150 bp sequencing, generating an estimated 2000 million clusters per lane). The raw Fastq files were trimmed for the presence of Illumina adapter sequences as described above.

Reference sequences were downloaded for each bacterial species. Trimmed reads were mapped to the reference genomes using bwa (identity>=90% and coverage>=60%). Reads mapped in proper pairs were extracted using samtools (-q 1 -f 2) ^120^. The number of mapped reads were summed up for each bacterial species in each sample, using custom scripts. To calculate scaled relative abundance, raw count data were first scaled by reference genome length, to account for the difference in genome length for different bacterial species. The scaled read count was divided by the sum count for each sample. To determine the presence and absence of each bacterial strain, reads were mapped from samples of each ancestral species to the reference genomes, the cutoff was determined by the number of reads that were randomly mapped to different genomes.

### Pathogen whole-genome sequencing

Whole genome sequencing was conducted on populations from passages 10 and 15 of evolved *S. aureus*. Populations consisted of 40 pooled clones per replicate. Each clone was cultured separately and standardised to the same optical density before being pooled in equal volumes. Pooled populations were then pelleted and stored at -20°C until used for DNA extractions. Simultaneously, two individual clones were sequenced per population.

Frozen pellets were resuspended in 200µl PBS for DNA extraction. Extractions were conducted using the High Pure PCR Template Preparation Kit (Roche) as per the manufacturers protocol with two changes: 10µg lysostaphin was added to lyse the cells for 30 mins instead of lysozyme and immediately following this samples were digested with 0.5µg RNase A for 15 mins at room temperature.

Sequencing was conducted by the Centre for Genomic Research at the University of Liverpool. Extracted DNA was prepared for Illumina short-read sequencing using the NEBNext Ultra II FS Kit on the Mosquito platform, using a 1/10 reduced volume protocol. DNA was sequenced by Illumina NovaSeq using SP chemistry (Paired-end, 2x150 bp sequencing, generating an estimated 325 million clusters per lane).

The raw Fastq files were trimmed for the presence of Illumina adapter sequences using Cutadapt version 1.2.1 ^112^. The option -O 3 was used, so the 3’ end of any reads which match the adapter sequence for 3 bp. or more are trimmed. The reads were further trimmed using Sickle version 1.200 with a minimum window quality score of 20. Reads shorter than 15 bp. after trimming were removed.

Reads were further trimmed using fastp ^121^. Variant calling was conducted using the breseq pipeline (v. 0.36.0) ^122^. Reads were mapped to the NCBI reference sequence NC_002953.3 (*Staphylococcus aureus* subsp. *aureus* MSSA476). gdtools was used to generate a comparison table of the SNPs in each sample and a spreadsheet detailing the type and number of mutations in each sample.

Euclidean genetic distance, both between replicates and of each replicate from the ancestor, was calculated using the dist() function in R.

Negative frequency dependent selection was calculated by regressing the change in pathogen variant frequencies (%) from passages 10-15 with the observed frequency (%) at passage 10, following the methods used by Koskella and Lively ^56,57^. Grey shading around the regression line represents 95% confidence intervals.

We calculated the nucleotide diversity across the entire pathogen genome for each replicate population to quantify the genetic diversity that arose from the ancestral clone. Nucleotide diversity, or pi, is the average number of nucleotide differences between all possible pairs of individuals in the population. We used the software PoPoolation^123^ to calculate nucleotide diversity (Tajima’s pi) using a sliding window analysis across the entire genome of evolved S. aureus. Each window size was 500bp with a step-size of 250bp. We calculated the mean of all windows with a positive value within each population, then the mean of all populations within each treatment across time.

### Pathogen qRT-PCR

Worms were colonised with either evolved or ancestral microbiota then infected with *S. aureus*, as described above. Only four replicates were assayed from each group for this experiment – the four most virulent from the pathogen that evolved alone and the four least virulent from the no-host control. After 12 hrs incubation on infection/microbiota plates (12hrs chosen for optimal gene expression), worms were washed off the plates with 3ml M9-Tx and left to settle at the bottom of a 15ml centrifuge tube. Worms were aspirated from 15ml tubes in 150µl volumes and transferred to cut-off 1000μl filter tips within 1.5ml tubes. These tubes were centrifuged for 2 mins at 290 x g, worms were then resuspended in 600µl RNA lysis buffer (Zymo) and transferred to ZR bead bashing tubes. Worms were lysed with a Disruptor Genie (full speed for 2 mins), then centrifuged for 5 mins at 16,000 x g. RNA was extracted from these lysed samples using the Zymo Quick RNA Miniprep kit, as per the manufacturers protocol and stored at -80°C.

DNA was digested using the Turbo DNase digest kit, as per the manufacturers protocol. cDNA was synthesized from RNA using the AccuScript High-Fidelity First Strand cDNA Synthesis kit, as per the manufacturers protocol. 100ng RNA was used for all cDNA synthesis reactions, to standardize the amount of RNA across samples. qPCR was conducted using the Luna qPCR Master Mix (NEB), with two technical replicates per cDNA sample. Briefly, 2µl cDNA was added to a 20µl reaction mix, containing 0.25µM final concentration of each primer. RNAIII primers (FW: AGCATGTAAGCTATCGTAAACAAC and RV: TTCAATCTATTTTTGGGGATG) were used as the target gene and gyrB primers (FW: CAGCGTTAGATGTAGCAAGC and RV: CCGATTCCTGTACCAAATGC) were used as the housekeeping gene. Expression of RNAIII was calculated relative to the housekeeping gene using the 2^-ΔΔCT^ method ^124^. For the evolved pathogen qRT-PCR, one outlier was removed at the data analysis stage, leaving three replicates per group in total.

### Genome-scale metabolic modelling

To explore the potential nutritional overlap and metabolic interactions between each microbiota species and the pathogen, we reconstructed genome-scale metabolic models (GEMs) for each strain. For each strain, its annotated protein sequences were downloaded from NCBI, and were used for GEM reconstruction based on the top-down carving approach of curated “universal models” using CarveMe ^125^. Pairwise potential cross-feeding interactions were evaluated using SMETANA scores, calculated using smetana (with 100 permutations) . SMETANA scores can range from 0 (complete independence) to 1 (essentiality) for each metabolite. For each bacterial pair, Metabolic Interaction Potential (MIP) and Metabolic Resource Overlap (MRO) were calculated using smetana. MIP calculates how many metabolites two species can share or exchange to decrease their dependency on external resources. It can be used to evaluate microbial cooperative metabolism. MIP and MRO estimate microbial interactions at their theoretical limit, regardless of the external environment.

### Statistics

All statistical analysis was conducted in R v4.2.2. Between 2-4 technical replicates were conducted for all assays and unless stated otherwise technical replicates were treated as independent data points. All data were tested for normality using the Shapiro-Wilk normality test. Normally distributed data was tested for statistical significance using parametric tests, including Welch Two-sample T-tests for datasets with two groups and ANOVA with post-hoc TukeyHSD tests for datasets with more than two groups. Non-normally distributed data was tested for statistical significance using non-parametric tests, including Wilcoxon rank sum tests for datasets with two groups and Kruskal-Wallis tests with post-hoc Dunn tests for datasets with more than two groups. Results were considered to be statistically significant when p<=0.05. P values were corrected for multiple comparisons using the Benjamini-Hochberg method. In cases when it was not relevant to compare all groups with each other, data was tested for statistical significance using a linear regression, and the ‘emmeans’ package was used to specify groups for post-hoc analysis.

Plots were made using the ‘ggplot2’ package in R^127^. Normally distributed data was plotted with the mean and standard error of the mean indicated on graphs. Non-normally distributed data was plotted with boxplots overlaying the datapoints to show the median and interquartile ranges. Whiskers of the boxplots represent 1.5 x interquartile range +/- third and first quartiles respectively. Points outside the whiskers are considered outliers.

## Supporting information

Supplementary file 1 - smetana scores

## Acknowledgements

We thank B. Samuel for sequencing data analysis support and H. Schulenburg for advice on experimental design.

## Author contributions

EJS and KCK conceived and designed the experiment. EJS conducted the experiments, with assistance from TEH and STEG. EJS conducted data analysis. EJS, JDL, and GCD conducted genomic analysis with guidance from SP. JDL conducted nutrient modelling. EJS and KCK wrote the initial draft with input from all authors.

## Data and materials availability

Phenotypic data is provided as an Excel file alongside the submitted manuscript. Genomic data can be viewed by reviewers at this link https://dataview.ncbi.nlm.nih.gov/object/PRJNA1055888?reviewer=ea120ks466cf8njtk6s0kphppu, and will be made publicly available upon publication.

## References

1. Ford, S. A. & King, K. C. Harnessing the Power of Defensive Microbes: Evolutionary Implications in Nature and Disease Control. PLOS Pathogens 12, e1005465 (2016).

2. Hoang, K. L. & King, K. C. Symbiosis: Partners in crime. Current Biology 32, R1018– R1020 (2022).

3. Brown, S. P., Le Chat, L. & Taddei, F. Evolution of virulence: triggering host inflammation allows invading pathogens to exclude competitors. Ecol Lett 11, 44–51 (2008).

4. Brown, S. P., Fredrik Inglis, R. & Taddei, F. Evolutionary ecology of microbial wars: within-host competition and (incidental) virulence. Evol Appl 2, 32–39 (2009).

5. Kamada, N., Chen, G. Y., Inohara, N. & Núñez, G. Control of Pathogens and Pathobionts by the Gut Microbiota. Nat Immunol 14, 685–690 (2013).

6. Guittar, J., Koffel, T., Shade, A., Klausmeier, C. A. & Litchman, E. Resource Competition and Host Feedbacks Underlie Regime Shifts in Gut Microbiota. The American Naturalist 198, 1–12 (2021).

7. Ford, S. A., Kao, D., Williams, D. & King, K. C. Microbe-mediated host defence drives the evolution of reduced pathogen virulence. Nature Communications 7, 1–9 (2016).

8. Tao, W. et al. Analysis of the intestinal microbiota in COVID-19 patients and its correlation with the inflammatory factor IL-18. Medicine in Microecology 5, 100023 (2020).

9. Houlden, A. et al. Chronic Trichuris muris Infection in C57BL/6 Mice Causes Significant Changes in Host Microbiota and Metabolome: Effects Reversed by Pathogen Clearance. PLOS ONE 10, e0125945 (2015).

10. Pharaon, J. & Bauch, C. T. The influence of social behaviour on competition between virulent pathogen strains. Journal of Theoretical Biology 455, 47–53 (2018).

11. López-Villavicencio, M. et al. COMPETITION, COOPERATION AMONG KIN, AND VIRULENCE IN MULTIPLE INFECTIONS. Evolution 65, 1357–1366 (2011).

12. Stevens, E. J., Bates, K. A. & King, K. C. Host microbiota can facilitate pathogen infection. PLOS Pathogens 17, e1009514 (2021).

13. Butel, M.-J. Probiotics, gut microbiota and health. Médecine et Maladies Infectieuses 44, 1–8 (2014).

14. Chua, K. J., Kwok, W. C., Aggarwal, N., Sun, T. & Chang, M. W. Designer probiotics for the prevention and treatment of human diseases. Current Opinion in Chemical Biology 40, 8–16 (2017).

15. Antwis, R. E. & Harrison, X. A. Probiotic consortia are not uniformly effective against different amphibian chytrid pathogen isolates. Molecular Ecology 27, 577–589 (2018).

16. Brugman, S. et al. A Comparative Review on Microbiota Manipulation: Lessons From Fish, Plants, Livestock, and Human Research. Front Nutr 5, 80 (2018).

17. McNally, L., Vale, P. F. & Brown, S. P. Microbiome engineering could select for more virulent pathogens. bioRxiv 027854 (2015) doi:10.1101/027854.

18. Nowak, M. A. & May, R. M. Superinfection and the evolution of parasite virulence. Proceedings of the Royal Society of London. Series B: Biological Sciences 255, 81–89 (1994).

19. Frank, S. A. Models of Parasite Virulence. The Quarterly Review of Biology 71, 37–78 (1996).

20. Barreto, H. C. & Gordo, I. Intrahost evolution of the gut microbiota. Nat Rev Microbiol 21, 590–603 (2023).

21. Freter, R., Brickner, H., Fekete, J., Vickerman, M. M. & Carey, K. E. Survival and implantation of Escherichia coli in the intestinal tract. Infect Immun 39, 686–703 (1983).

22. Levin, B. R. & Bull, J. J. Short-sighted evolution and the virulence of pathogenic microorganisms. Trends Microbiol 2, 76–81 (1994).

23. Barroso-Batista, J. et al. Specific Eco-evolutionary Contexts in the Mouse Gut Reveal Escherichia coli Metabolic Versatility. Current Biology 30, 1049–1062.e7 (2020).

24. Barroso-Batista, J. et al. The First Steps of Adaptation of Escherichia coli to the Gut Are Dominated by Soft Sweeps. PLOS Genetics 10, e1004182 (2014).

25. King, K. C. et al. Rapid evolution of microbe-mediated protection against pathogens in a worm host. The ISME Journal 10, 1915–1924 (2016).

26. Hall, J. P. J., Harrison, E. & Brockhurst, M. A. Competitive species interactions constrain abiotic adaptation in a bacterial soil community. Evolution Letters 2, 580–589 (2018).

27. Johansson, J. EVOLUTIONARY RESPONSES TO ENVIRONMENTAL CHANGES: HOW DOES COMPETITION AFFECT ADAPTATION? Evolution 62, 421–435 (2008).

28. Drew, G. C., Stevens, E. J. & King, K. C. Microbial evolution and transitions along the parasite–mutualist continuum. Nat Rev Microbiol 19, 623–638 (2021).

29. Montalvo-Katz, S., Huang, H., Appel, M. D., Berg, M. & Shapira, M. Association with Soil Bacteria Enhances p38-Dependent Infection Resistance in Caenorhabditis elegans. Infection and Immunity 81, 514–520 (2013).

30. Rossouw, W. & Korsten, L. Cultivable microbiome of fresh white button mushrooms. Lett Appl Microbiol 64, 164–170 (2017).

31. Grewal, P. S. Relative contribution of nematodes (Caenorhabditis elegans) and bacteria towards the disruption of flushing patterns and losses in yield and quality of mushrooms (Agaricus bisporus). Annals of Applied Biology 119, 483–499 (1991).

32. Mrochen, D. M. et al. Wild rodents and shrews are natural hosts of Staphylococcus aureus. International Journal of Medical Microbiology 308, 590–597 (2018).

33. Peton, V. & Le Loir, Y. Staphylococcus aureus in veterinary medicine. Infect Genet Evol 21, 602–615 (2014).

34. Schaumburg, F. et al. Drug-resistant human Staphylococcus aureus in sanctuary apes pose a threat to endangered wild ape populations. Am J Primatol 74, 1071–1075 (2012).

35. Frank, D. N. et al. The Human Nasal Microbiota and Staphylococcus aureus Carriage. PLOS ONE 5, e10598 (2010).

36. Lina, G. et al. Bacterial Competition for Human Nasal Cavity Colonization: Role of Staphylococcal agr Alleles. Applied and Environmental Microbiology 69, 18–23 (2003).

37. Margolis, E., Yates, A. & Levin, B. R. The ecology of nasal colonization of Streptococcus pneumoniae, Haemophilus influenzae and Staphylococcus aureus: the role of competition and interactions with host’s immune response. BMC Microbiol 10, 59 (2010).

38. Rudkin, J. K., McLoughlin, R. M., Preston, A. & Massey, R. C. Bacterial toxins: Offensive, defensive, or something else altogether? PLoS Pathog 13, (2017).

39. Irazoqui, J. E. et al. Distinct Pathogenesis and Host Responses during Infection of C. elegans by P. aeruginosa and S. aureus. PLOS Pathogens 6, e1000982 (2010).

40. Sifri, C. D., Begun, J., Ausubel, F. M. & Calderwood, S. B. Caenorhabditis elegans as a Model Host for Staphylococcus aureus Pathogenesis. Infection and Immunity 71, 2208– 2217 (2003).

41. Thompson, T. A. & Brown, P. D. Association between the agr locus and the presence of virulence genes and pathogenesis in Staphylococcus aureus using a Caenorhabditis elegans model. International Journal of Infectious Diseases 54, 72–76 (2017).

42. Dirksen, P. et al. CeMbio - The Caenorhabditis elegans Microbiome Resource. G3 Genes|Genomes|Genetics 10, 3025–3039 (2020).

43. Recsei, P. et al. Regulation of exoprotein gene expression in Staphylococcus aureus by agr. Mol Gen Genet 202, 58–61 (1986).

44. Abdelnour, A., Arvidson, S., Bremell, T., Rydén, C. & Tarkowski, A. The accessory gene regulator (agr) controls Staphylococcus aureus virulence in a murine arthritis model. Infect Immun 61, 3879–3885 (1993).

45. Traber, K. E. et al. agr function in clinical Staphylococcus aureus isolates. Microbiology (Reading) 154, 2265–2274 (2008).

46. Novick, R. P. Autoinduction and signal transduction in the regulation of staphylococcal virulence. Molecular Microbiology 48, 1429–1449 (2003).

47. Novick, R. P. et al. Theagr P2 operon: An autocatalytic sensory transduction system inStaphylococcus aureus. Molec. Gen. Genet. 248, 446–458 (1995).

48. Graf, A. C. et al. Virulence Factors Produced by Staphylococcus aureus Biofilms Have a Moonlighting Function Contributing to Biofilm Integrity *[S]. Molecular & Cellular Proteomics 18, 1036–1053 (2019).

49. Cheung, G. Y. C., Bae, J. S. & Otto, M. Pathogenicity and virulence of Staphylococcus aureus. Virulence 12, 547–569 (2021).

50. McCarthy, H. et al. Methicillin resistance and the biofilm phenotype in Staphylococcus aureus. Frontiers in Cellular and Infection Microbiology 5, (2015).

51. Begun, J. et al. Staphylococcal Biofilm Exopolysaccharide Protects against Caenorhabditis elegans Immune Defenses. PLOS Pathogens 3, e57 (2007).

52. Carver, T. J. et al. ACT: the Artemis comparison tool. Bioinformatics 21, 3422–3423 (2005).

53. Chen, H. et al. Polygenic virulence factors involved in pathogenesis of 1997 Hong Kong H5N1 influenza viruses in mice. Virus Res 128, 159–163 (2007).

54. Caseys, C., et al. Quantitative interactions: the disease outcome of Botrytis cinerea across the plant kingdom. G3 Genes|Genomes|Genetics 11, jkab175 (2021).

55. Le Clec’h, W. et al. Genetic architecture of transmission stage production and virulence in schistosome parasites. Virulence 12, 1508–1526 (2021).

56. Koskella, B. & Lively, C. M. Evidence for Negative Frequency-Dependent Selection during Experimental Coevolution of a Freshwater Snail and a Sterilizing Trematode. Evolution 63, 2213–2221 (2009).

57. Betts, A., Gray, C., Zelek, M., MacLean, R. C. & King, K. C. High parasite diversity accelerates host adaptation and diversification. Science 360, 907–911 (2018).

58. Christie, M. R. & McNickle, G. G. Negative frequency dependent selection unites ecology and evolution. Ecology and Evolution 13, e10327 (2023).

59. Brisson, D. Negative Frequency-Dependent Selection Is Frequently Confounding. Frontiers in Ecology and Evolution 6, (2018).

60. Ghoul, M. & Mitri, S. The Ecology and Evolution of Microbial Competition. Trends in Microbiology 24, 833–845 (2016).

61. Hibbing, M. E., Fuqua, C., Parsek, M. R. & Peterson, S. B. Bacterial competition: surviving and thriving in the microbial jungle. Nat Rev Microbiol 8, 15–25 (2010).

62. May, R. M. & Nowak, M. A. Coinfection and the evolution of parasite virulence. Proceedings of the Royal Society of London. Series B: Biological Sciences 261, 209–215 (1997).

63. Chang, J. Y. et al. Decreased diversity of the fecal Microbiome in recurrent Clostridium difficile-associated diarrhea. J Infect Dis 197, 435–438 (2008).

64. Rabajante, J. F. et al. Red Queen dynamics in multi-host and multi-parasite interaction system. Sci Rep 5, 10004 (2015).

65. Mack, K. M. L., Eppinga, M. B. & Bever, J. D. Plant-soil feedbacks promote coexistence and resilience in multi-species communities. PLOS ONE 14, e0211572 (2019).

66. Feil, E. J. et al. How clonal is Staphylococcus aureus? J Bacteriol 185, 3307–3316 (2003).

67. Paterson, S. et al. Antagonistic coevolution accelerates molecular evolution. Nature 464, 275–278 (2010).

68. Barreto, H. C., Abreu, B. & Gordo, I. Fluctuating selection on bacterial iron regulation in the mammalian gut. Curr Biol 32, 3261–3275.e4 (2022).

69. Rafaluk, C., Jansen, G., Schulenburg, H. & Joop, G. When experimental selection for virulence leads to loss of virulence. Trends in Parasitology 31, 426–434 (2015).

70. Beenken, K. E. et al. Epistatic Relationships between sarA and agr in Staphylococcus aureus Biofilm Formation. PLOS ONE 5, e10790 (2010).

71. Joshua, G. W. P. et al. A Caenorhabditis elegans model of Yersinia infection: biofilm formation on a biotic surface. Microbiology 149, 3221–3229 (2003).

72. Gillaspy, A. F. et al. Role of the accessory gene regulator (agr) in pathogenesis of staphylococcal osteomyelitis. Infect Immun 63, 3373–3380 (1995).

73. Kielian, T., Cheung, A. & Hickey, W. F. Diminished Virulence of an Alpha-Toxin Mutant ofStaphylococcus aureus in Experimental Brain Abscesses. Infection and Immunity 69, 6902–6911 (2001).

74. Cheung, A. L. et al. Diminished virulence of a sar-/agr- mutant of Staphylococcus aureus in the rabbit model of endocarditis. J Clin Invest 94, 1815–1822 (1994).

75. Pollitt, E. J. G., West, S. A., Crusz, S. A., Burton-Chellew, M. N. & Diggle, S. P. Cooperation, Quorum Sensing, and Evolution of Virulence in Staphylococcus aureus. Infect. Immun. 82, 1045–1051 (2014).

76. Cafiso, V. et al. agr-Genotyping and transcriptional analysis of biofilm-producing Staphylococcus aureus. FEMS Immunology & Medical Microbiology 51, 220–227 (2007).

77. Lauderdale, K. J., Boles, B. R., Cheung, A. L. & Horswill, A. R. Interconnections between Sigma B, agr, and Proteolytic Activity in Staphylococcus aureus Biofilm Maturation. Infection and Immunity 77, 1623–1635 (2009).

78. Yarwood, J. M., Bartels, D. J., Volper, E. M. & Greenberg, E. P. Quorum Sensing in Staphylococcus aureus Biofilms. Journal of Bacteriology 186, 1838–1850 (2004).

79. Pham, T. A. N. & Lawley, T. D. Emerging insights on intestinal dysbiosis during bacterial infections. Current Opinion in Microbiology 17, 67–74 (2014).

80. de Steenhuijsen Piters, W. A. A., et al. Dysbiosis of upper respiratory tract microbiota in elderly pneumonia patients. ISME J 10, 97–108 (2016).

81. Fletcher, J. R. et al. Clostridioides difficile exploits toxin-mediated inflammation to alter the host nutritional landscape and exclude competitors from the gut microbiota. Nat Commun 12, 462 (2021).

82. Zaneveld, J. R., McMinds, R. & Vega Thurber, R. Stress and stability: applying the Anna Karenina principle to animal microbiomes. Nature Microbiology 2, 1–8 (2017).

83. Ma, Z. S. Testing the Anna Karenina Principle in Human Microbiome-Associated Diseases. iScience 23, 101007 (2020).

84. Foster, K. R., Schluter, J., Coyte, K. Z. & Rakoff-Nahoum, S. The evolution of the host microbiome as an ecosystem on a leash. Nature 548, 43–51 (2017).

85. Beatty, J. K. et al. Giardia duodenalis induces pathogenic dysbiosis of human intestinal microbiota biofilms. International Journal for Parasitology 47, 311–326 (2017).

86. Wei, Z. et al. Ralstonia solanacearum pathogen disrupts bacterial rhizosphere microbiome during an invasion. Soil Biology and Biochemistry 118, 8–17 (2018).

87. Lowder, B. V. et al. Recent human-to-poultry host jump, adaptation, and pandemic spread of Staphylococcus aureus. Proceedings of the National Academy of Sciences 106, 19545–19550 (2009).

88. Spoor, L. E. et al. Livestock Origin for a Human Pandemic Clone of Community- Associated Methicillin-Resistant Staphylococcus aureus. mBio 4, 10.1128/mbio.00356-13 (2013).

89. Senghore, M. et al. Transmission of Staphylococcus aureus from Humans to Green Monkeys in The Gambia as Revealed by Whole-Genome Sequencing. Applied and Environmental Microbiology 82, 5910–5917 (2016).

90. Vincze, S. et al. Alarming Proportions of Methicillin-Resistant Staphylococcus aureus (MRSA) in Wound Samples from Companion Animals, Germany 2010–2012. PLOS ONE 9, e85656 (2014).

91. Krismer, B., Weidenmaier, C., Zipperer, A. & Peschel, A. The commensal lifestyle of Staphylococcus aureus and its interactions with the nasal microbiota. Nat Rev Microbiol 15, 675–687 (2017).

92. Liu, C. M. et al. Staphylococcus aureus and the ecology of the nasal microbiome. Science Advances 1, e1400216 (2015).

93. Kastman, E. K. et al. Biotic Interactions Shape the Ecological Distributions of Staphylococcus Species. mBio 7, 10.1128/mbio.01157-16 (2016).

94. Yan, M. et al. Nasal Microenvironments and Interspecific Interactions Influence Nasal Microbiota Complexity and S. aureus Carriage. Cell Host & Microbe 14, 631–640 (2013).

95. Power, A. G. & Mitchell, C. E. Pathogen Spillover in Disease Epidemics. The American Naturalist 164, S79–S89 (2004).

96. Borremans, B., Faust, C., Manlove, K. R., Sokolow, S. H. & Lloyd-Smith, J. O. Cross- species pathogen spillover across ecosystem boundaries: mechanisms and theory. Philosophical Transactions of the Royal Society B: Biological Sciences 374, 20180344 (2019).

97. Cohen, J. M., Sauer, E. L., Santiago, O., Spencer, S. & Rohr, J. R. Divergent impacts of warming weather on wildlife disease risk across climates. Science 370, eabb1702 (2020).

98. Gibb, R. et al. Zoonotic host diversity increases in human-dominated ecosystems. Nature 584, 398–402 (2020).

99. Agrawal, A. & Lively, C. M. Infection genetics: gene-for-gene versus matching- alleles models and all points in between.

100. Aleman, F. D. D. & Valenzano, D. R. Microbiome evolution during host aging. PLOS Pathogens 15, e1007727 (2019).

101. Dapa, T., Ramiro, R. S., Pedro, M. F., Gordo, I. & Xavier, K. B. Diet leaves a genetic signature in a keystone member of the gut microbiota. Cell Host Microbe 30, 183–199.e10 (2022).

102. Aballay, A. & Ausubel, F. M. Caenorhabditis elegans as a host for the study of host– pathogen interactions. Current Opinion in Microbiology 5, 97–101 (2002).

103. Clark, L. C. & Hodgkin, J. Commensals, probiotics and pathogens in the Caenorhabditis elegans model. Cellular Microbiology 16, 27–38 (2014).

104. Hilbi, H., Weber, S. S., Ragaz, C., Nyfeler, Y. & Urwyler, S. Environmental predators as models for bacterial pathogenesis. Environmental Microbiology 9, 563–575 (2007).

105. Laaberki, M.-H. & Dworkin, J. Role of Spore Coat Proteins in the Resistance of Bacillus subtilis Spores to Caenorhabditis elegans Predation. Journal of Bacteriology 190, 6197– 6203 (2008).

106. Lee, J.-H. et al. Indole-associated predator–prey interactions between the nematode Caenorhabditis elegans and bacteria. Environmental Microbiology 19, 1776–1790 (2017).

107. Burlinson, P. et al. Pseudomonas fluorescens NZI7 repels grazing by C. elegans, a natural predator. ISME J 7, 1126–1138 (2013).

108. Marsh, E. K. & May, R. C. Caenorhabditis elegans, a Model Organism for Investigating Immunity. Applied and Environmental Microbiology 78, 2075–2081 (2012).

109. Holden, M. T. G. et al. Complete genomes of two clinical Staphylococcus aureus strains: Evidence for the rapid evolution of virulence and drug resistance. Proceedings of the National Academy of Sciences 101, 9786–9791 (2004).

110. Wingett, S. W. & Andrews, S. FastQ Screen: A tool for multi-genome mapping and quality control. F1000Res 7, 1338 (2018).

111. Ewels, P., Magnusson, M., Lundin, S. & Käller, M. MultiQC: summarize analysis results for multiple tools and samples in a single report. Bioinformatics 32, 3047–3048 (2016).

112. Martin, M. Cutadapt removes adapter sequences from high-throughput sequencing reads. EMBnet.journal 17, 10–12 (2011).

113. Rognes, T., Flouri, T., Nichols, B., Quince, C. & Mahé, F. VSEARCH: a versatile open source tool for metagenomics. PeerJ 4, e2584 (2016).

114. Amir, A. et al. Deblur Rapidly Resolves Single-Nucleotide Community Sequence Patterns. mSystems 2, e00191–16 (2017).

115. Bolyen, E. et al. Reproducible, interactive, scalable and extensible microbiome data science using QIIME 2. Nat Biotechnol 37, 852–857 (2019).

116. Oksanen, et al. _vegan: Community Ecology Package_. R package version 2.6–4. (2022).

117. Madeira, F. et al. Search and sequence analysis tools services from EMBL-EBI in 2022. Nucleic Acids Res 50, W276–W279 (2022).

118. Chernomor, O., von Haeseler, A. & Minh, B. Q. Terrace Aware Data Structure for Phylogenomic Inference from Supermatrices. Systematic Biology 65, 997–1008 (2016).

119. Kalyaanamoorthy, S., Minh, B. Q., Wong, T. K. F., von Haeseler, A. & Jermiin, L. S. ModelFinder: fast model selection for accurate phylogenetic estimates. Nat Methods 14, 587–589 (2017).

120. Danecek, P. et al. Twelve years of SAMtools and BCFtools. GigaScience 10, giab008 (2021).

121. Chen, S., Zhou, Y., Chen, Y. & Gu, J. fastp: an ultra-fast all-in-one FASTQ preprocessor. Bioinformatics 34, i884–i890 (2018).

122. Deatherage, D. E. & Barrick, J. E. Identification of mutations in laboratory evolved microbes from next-generation sequencing data using breseq. Methods Mol Biol 1151, 165–188 (2014).

123. Kofler, R. et al. PoPoolation: A Toolbox for Population Genetic Analysis of Next Generation Sequencing Data from Pooled Individuals. PLOS ONE 6, e15925 (2011).

124. Schmittgen, T. D. & Livak, K. J. Analyzing real-time PCR data by the comparative CT method. Nat Protoc 3, 1101–1108 (2008).

125. Machado, D., Andrejev, S., Tramontano, M. & Patil, K. R. Fast automated reconstruction of genome-scale metabolic models for microbial species and communities. Nucleic Acids Res 46, 7542–7553 (2018).

126. Zelezniak, A. et al. Metabolic dependencies drive species co-occurrence in diverse microbial communities. Proceedings of the National Academy of Sciences 112, 6449– 6454 (2015).

127. Wickham, H. *Ggplot2: Elegant Graphics for Data Analysis*. (Springer International Publishing, Cham, 2016). doi:10.1007/978-3-319-24277-4.

