## Supplementary figures for "Within-host competition sparks pathogen molecular evolution and perpetual microbiota dysbiosis"

**Supplementary figures and tables**

**Supplementary figure 1. S1a)** No difference in in vivo RNAIII expression of the ancestral pathogen in the presence and absence of the ancestral microbiota (n=4, Welch Two-sample T test, p=0.22). **S1b)** In the presence of ancestral microbiota, there is no difference in host mortality from each evolved pathogen (n=23-24, Kruskal Wallis rank sum test, df=2, p=0.373). **S1c)** All evolved pathogens colonise *C. elegans* significantly better in the presence, compared to the absence, of ancestral microbiota (n=12-17, linear regression using log10 of raw data, f=8.594, df=5, p<0.0001, pairwise comparisons of each group in the presence vs absence of ancestral microbiota, using 'emmeans' package: No-host control p=0.023, Pathogen only p<0.0001, Pathogen + anc mbiota p=0.014) **S1d)** Colonisation of *C. elegans* by the ancestral microbiota is significantly reduced by the presence of all evolved pathogens compared to the microbiota only (n=3 for microbiota only control, n=16-18 for all other groups, linear regression using log10 of raw data, f=64.53, df=6, p<0.0001, pairwise comparisons between mbiota CFU/worm using 'emmeans' package: Microbiota only vs No-host control p=0.0006, Microbiota only vs Pathogen only p=0.025, Microbiota only vs Pathogen + anc mbiota p=0.005), and all evolved pathogens colonise *C. elegans* to a significantly greater extent than the microbiota (n=16-18, linear regression using log10 of raw data, f=64.53, df=6, p<0.0001, pairwise comparisons between pathogen CFU/worm and mbiota CFU/worm in each evolved group: all p<0.0001).

**Supplementary figure 2.** No significant differences in alpha diversity of the ancestral microbiota community in the presence of each evolved pathogen in vitro, based on 16S sequencing (n=6, Kruskal Wallis rank sum test for Observed diversity p=0.57, ANOVA for Shannon diversity p=0.06, for Simpson p=0.12 and for Evenness p=0.21) or in vivo (n=6, Kruskal Wallis rank sum test for Observed diversity p=0.57, for Simpson p=0.37 and for Evenness p=0.53, ANOVA for Shannon diversity p=0.65).

**Supplementary figure 3. S3a)** Composition data from 16S sequencing of in vivo samples. (PERMANOVA Bray Curtis pseudo-F=2.4583, p=0.047, pairwise comparison yields significant difference between 'Pathogen only' and 'No-host control', pseudo-F=3.155, p=0.051). **S3b)** Composition data from 16S sequencing of in vitro samples. No significant differences in community composition across evolved groups (PERMANOVA Bray-Curtis pseudo-F=1.5094, p=0.2176).

**Supplementary figure 4.** Boxplot showing dispersion of microbiota communities in presence of each evolved pathogen. No significant differences between evolved groups in vitro (Bray-Curtis f=0.0774, df=2, p=0.932). Signficant difference between 'Pathogen + anc mbiota' group and 'No-host control' in vivo (Bray-Curtis, f=13.18, df=1, p=0.015).

**Supplementary figure 5. S5a)** Significantly fewer non-synonymous mutations (n=12, ANOVA f=6.76, df=2, p=0.004, TukeyHSD comparison between No-host control and Pathogen only p=0.003) and indels (n=12, Kruskal Wallis rank sum test df=2, p=0.02, Dunn test comparison between No-host control and Pathogen only p=0.02) account for the difference in total number of mutations in clones of the 'Pathogen only' group. **S5b)** Genetic distance from the ancestor of each replicate population within each evolutionary group at generations 10 and 15. No significant differences between groups (n=6, ANOVA f=1.99, df=3, p=0.14). **S5c)** Nucleotide diversity averaged across genome for each treatment group. No significant differences in nucleotide diversity were found between treatments or within treatments over time (comparison of all four groups - P10 and P15 from Pathogen + anc mbiota group and Pathogen only group - using Kruskal-Wallis rank sum test df=3, p=0.4689). **S5d)** Mutation frequency in each replicate of passages 10 and 15 for each treatment. The frequency of mutations ranged from 5-100%. **S5e)** Nonsynonymous and synonymous mutation counts across passages 10 and 15 for each treatment. In both treatments, there were generally more nonsynonymous than synonymous SNPs at each time point. **S5f)** Histogram of nonsynonymous and synonymous mutations for passages 10 and 15 of both treatments. Each tick mark on the x-axis indicates the higher value of that bin. E.g., the tick mark at 0.2 is showing how many non/syn mutations are found between 0.1 and 0.2 frequency. **S5g & h)** Frequencies of nonsynonymous and synonymous mutations at passages 10 and 15 for each treatment. Across populations, there was variation in terms of the frequency of nonsynonymous vs. synonymous SNPs. For example, most mutations in pathogen only P15 population six were at < 25%, whereas populations one and three of the same treatment and time point had many mutations at > 50%. **S5i)** Distribution of nonsynonymous and synonymous mutations across pathogen genome, per treatment for passages 10 and 15. **S5j)** Mutation frequency at passages 10 and 15 for replicates of the pathogen that evolved with ancestral microbiota. **S5k)** Mutation frequency at passages 10 and 15 for replicates of the pathogen that evolved in isolation.

**Supplementary figure 6.** No difference in *in vivo* RNAIII expression of the ancestral pathogen in the presence of the microbiota that evolved alone and the no-host control evolved community (n=4, Welch Two-sample T test p=0.924).

**Supplementary figure 7.** When colonised with only the evolved microbiota lineages and no pathogen, the microbiota that evolved alone caused slightly higher mortality in *C. elegans*, which was significantly different from the Mbiota + anc pathogen, but not significantly different to the no-host control (n=17-18, Kruskal Wallis rank sum test, df=2, p=0.01, Dunn test comparisons: No-host control vs Mbiota only p=0.099, Mbiota only vs Mbiota + anc pathogen p=0.012, No-host control vs Mbiota + anc pathogen p=0.29).

**Supplementary figure 8.** Overview of newly created evolving group at passage 10, in which the evolved 'Microbiota only' community was introduced to duplicates of the 'Pathogen only' group. This new group evolved in parallel to the original 6 groups for the remaining 5 passages of the evolution experiment.

**Supplementary figure 9.** When the evolved 'Microbiota only' community was introduced to duplicates of the 'Pathogen only' group at passage 10, no significant difference in virulence was observed at passage 15 between the respective pathogens from these two groups (n=18, Kruskal Wallis rank sum test, df=1, p=0.11).

**S1a**

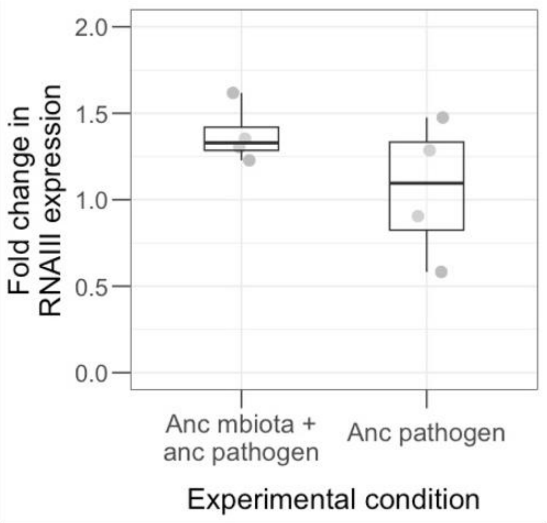


**S1b**

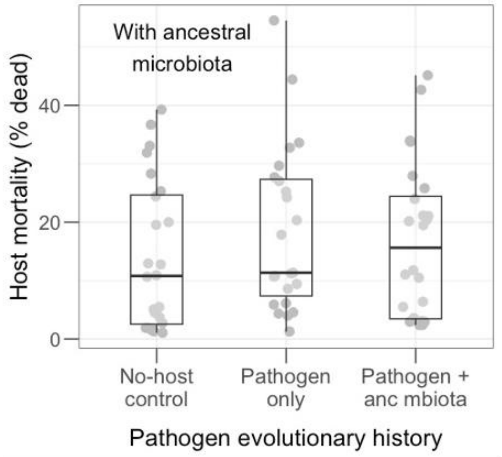


**S1c**

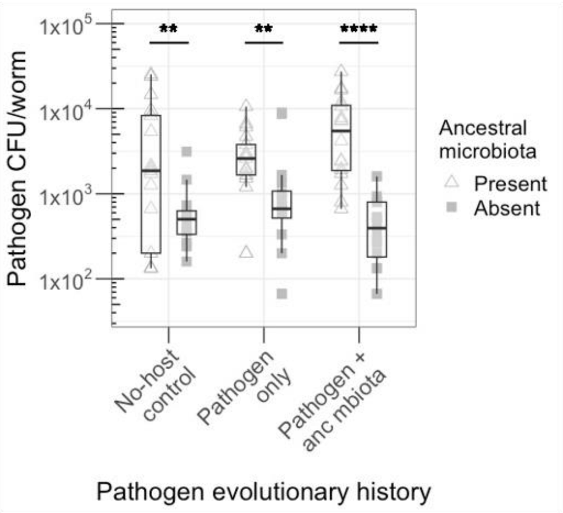


**S1d**

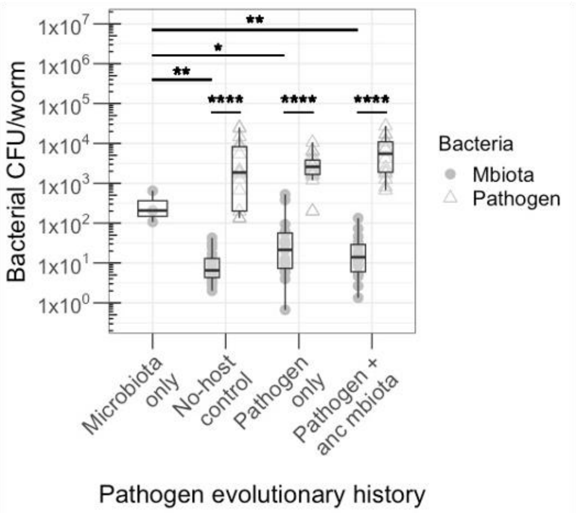

**S2**

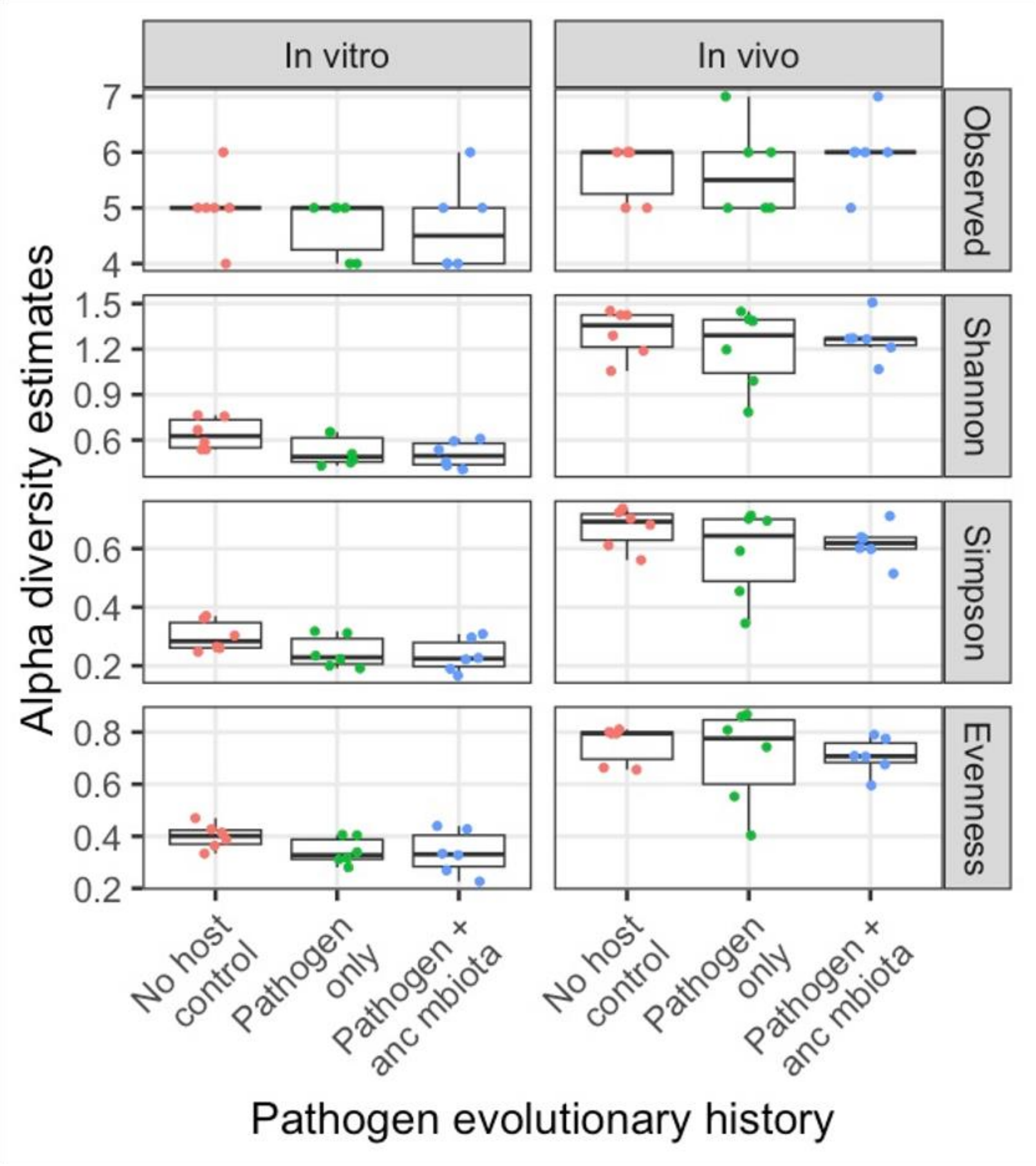

In vitro
In vivo
Observed
Shannon
Simpson
Evenness
Alpha diversity estimates
7
6
5
4
1.5
1.2
0.9
0.6
0.6
0.4
0.2
0.8
0.6
0.4
0.2
No host control
Pathogen only
Pathogen + anc mbiota
Pathogen evolutionary history

**S3a**

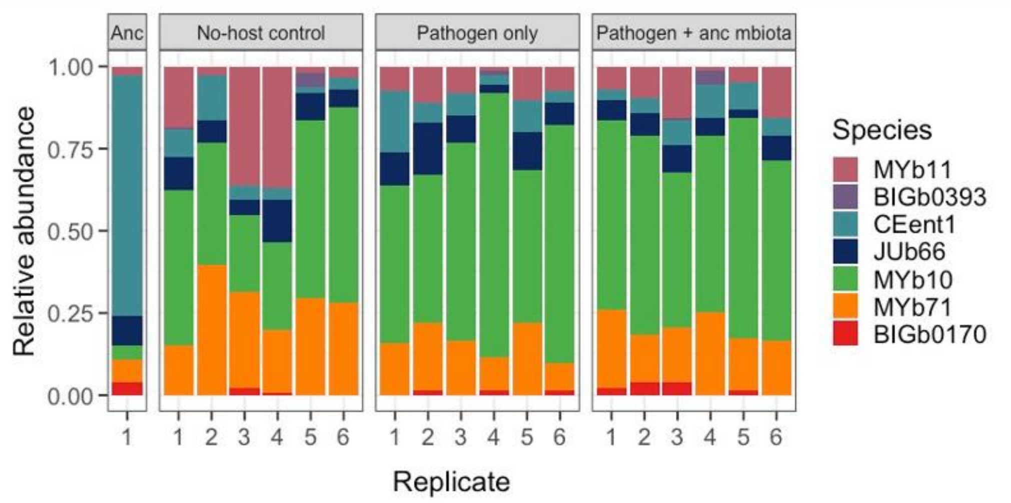
Anc
No-host control
Pathogen only
Pathogen + anc mbiota
Relative abundance
1.00
0.75
0.50
0.25
0.00
1
1 2 3 4 5 6
1 2 3 4 5 6
1 2 3 4 5 6
Replicate
Species
MYb11
BIGb0393
CEent1
JUb66
MYb10
MYb71
BIGb0170

**S3b**

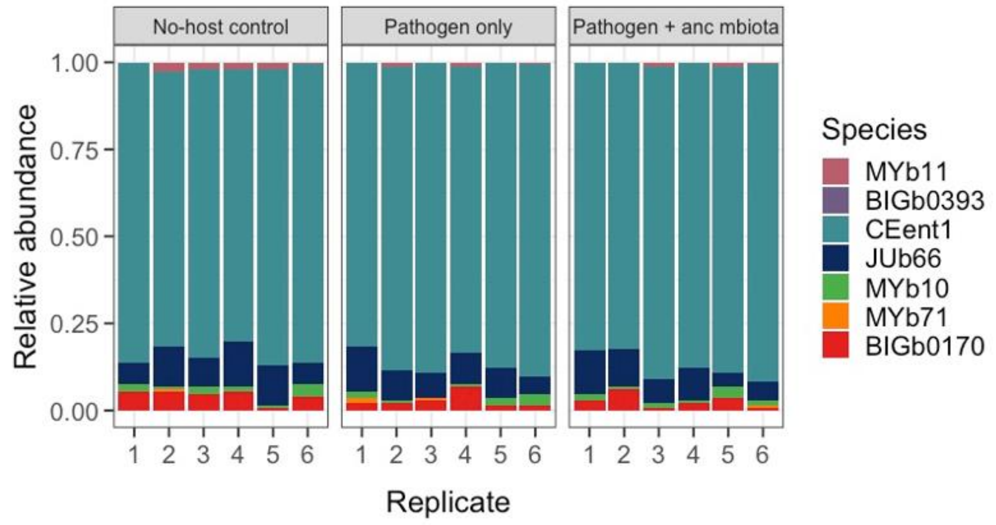
No-host control
Pathogen only
Pathogen + anc mbiota
Relative abundance
1.00
0.75
0.50
0.25
0.00
1 2 3 4 5 6
1 2 3 4 5 6
1 2 3 4 5 6
Replicate
Species
MYb11
BIGb0393
CEent1
JUb66
MYb10
MYb71
BIGb0170

**S4**

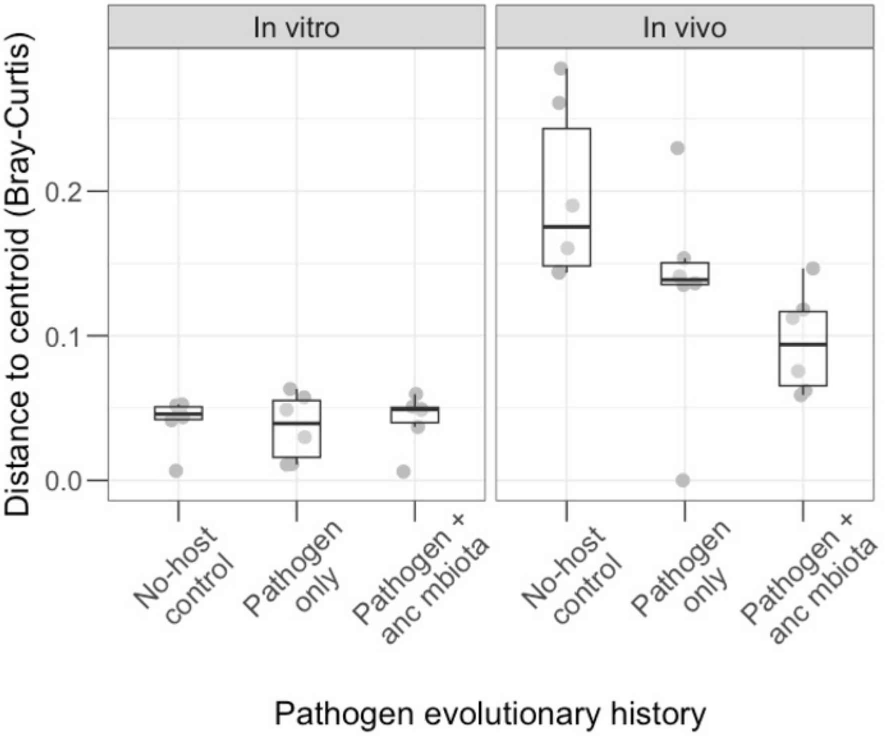
In vitro
In vivo
Distance to centroid (Bray-Curtis)
0.0
0.1
0.2
No-host control
Pathogen only
Pathogen + anc mbiota
No-host control
Pathogen only
Pathogen + anc mbiota
Pathogen evolutionary history

**S5a**

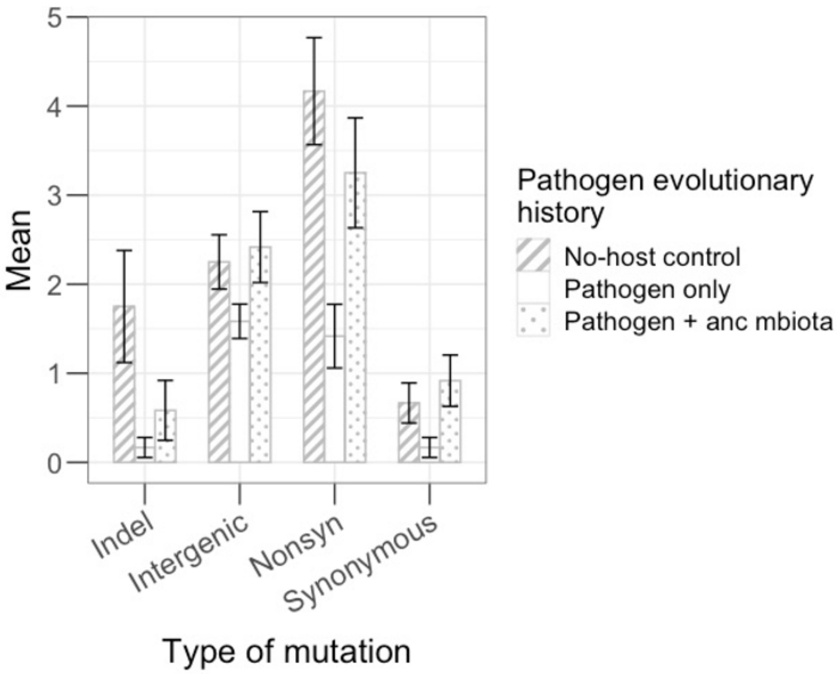

Mean
Type of mutation
Indel
Intergenic
Nonsyn
Synonymous
Pathogen evolutionary history
No-host control
Pathogen only
Pathogen + anc mbiota


**S5b**

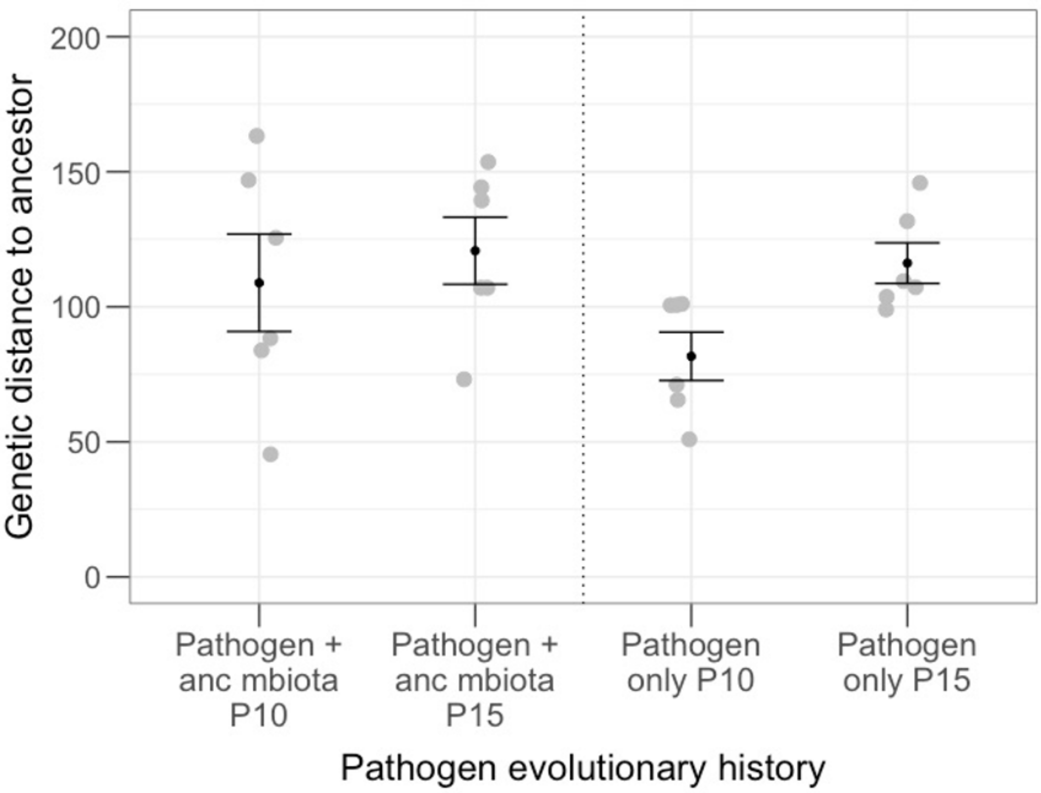

Genetic distance to ancestor
Pathogen + anc mbiota P10
Pathogen + anc mbiota P15
Pathogen only P10
Pathogen only P15
Pathogen evolutionary history

**S5c**

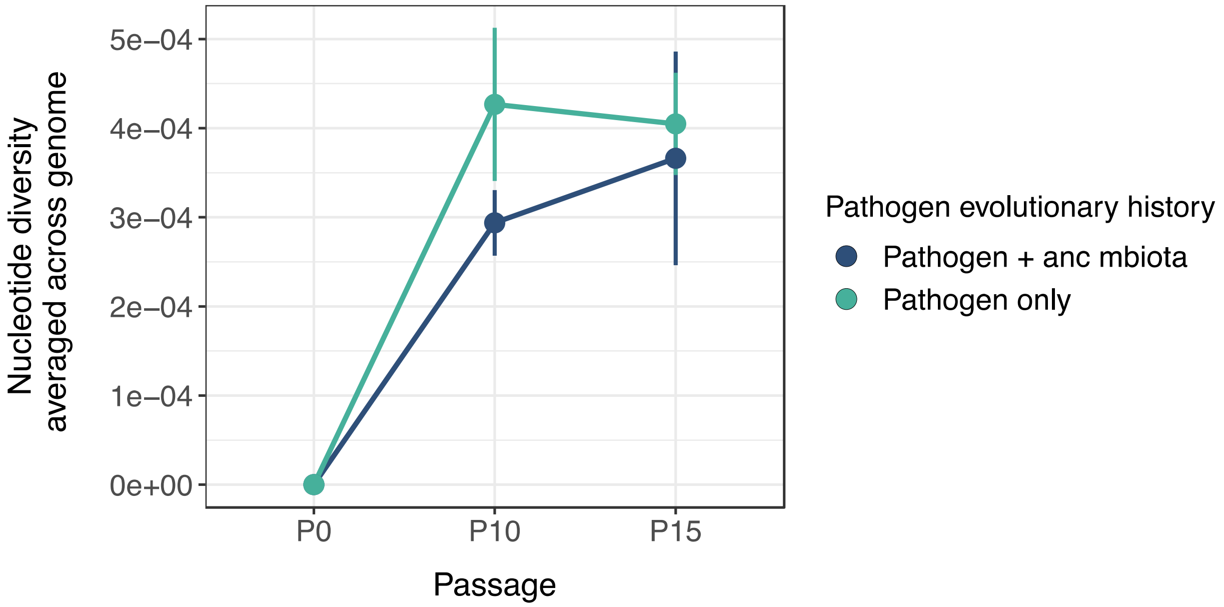
Nucleotide diversity
averaged across genome
5e−04
4e−04
3e−04
2e−04
1e−04
0e+00
P0
P10
P15
Passage
Pathogen evolutionary history
Pathogen + anc mbiota
Pathogen only

**S5d**

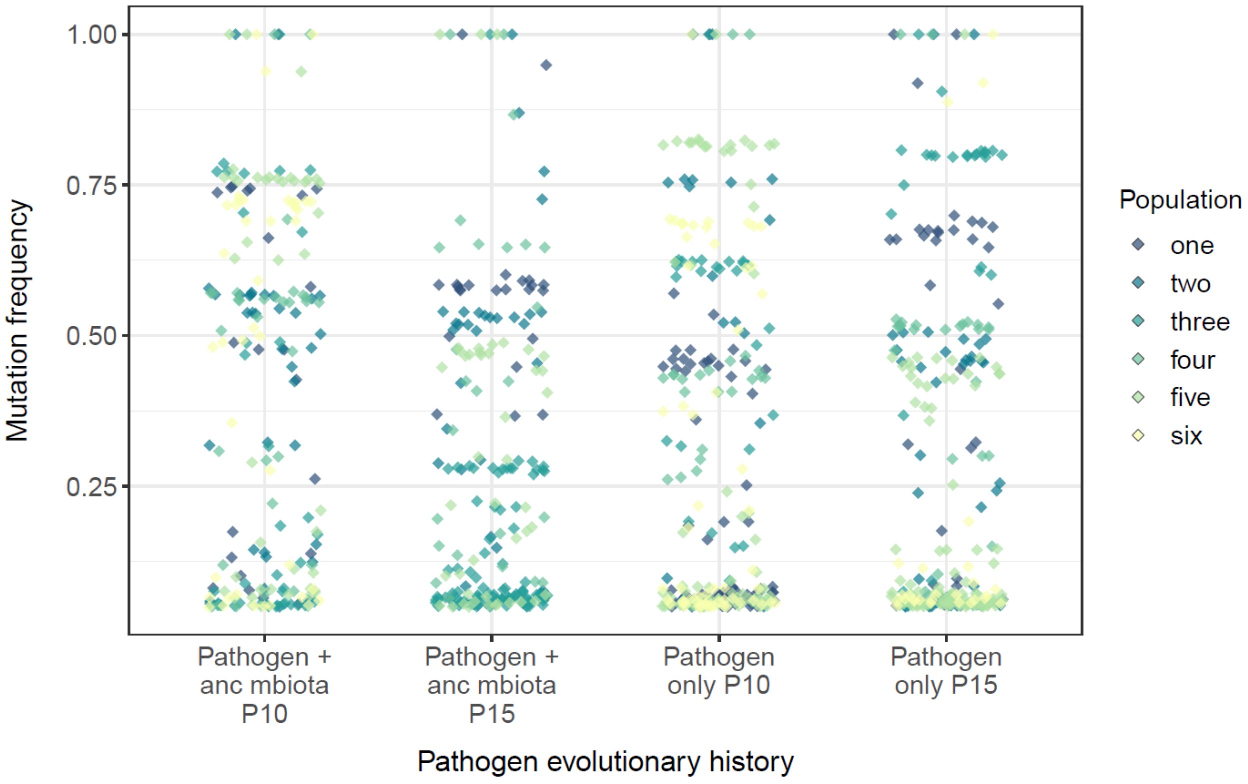
Mutation frequency
1.00
0.75
0.50
0.25
Pathogen + anc mbiota P10
Pathogen + anc mbiota P15
Pathogen only P10
Pathogen only P15
Pathogen evolutionary history
Population
one
two
three
four
five
six

**S5e**

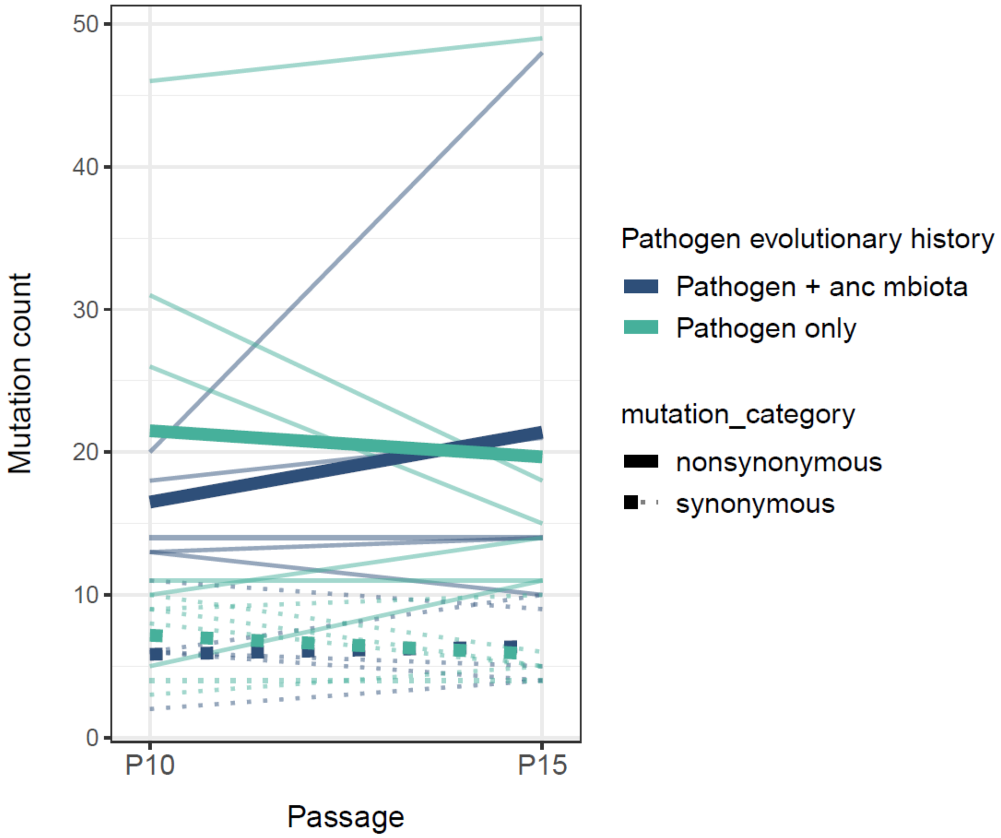
Mutation count
50
40
30
20
10
0
P10
P15
Passage
Pathogen evolutionary history
Pathogen + anc mbiota
Pathogen only
mutation_category
nonsynonymous
synonymous

**S5f**

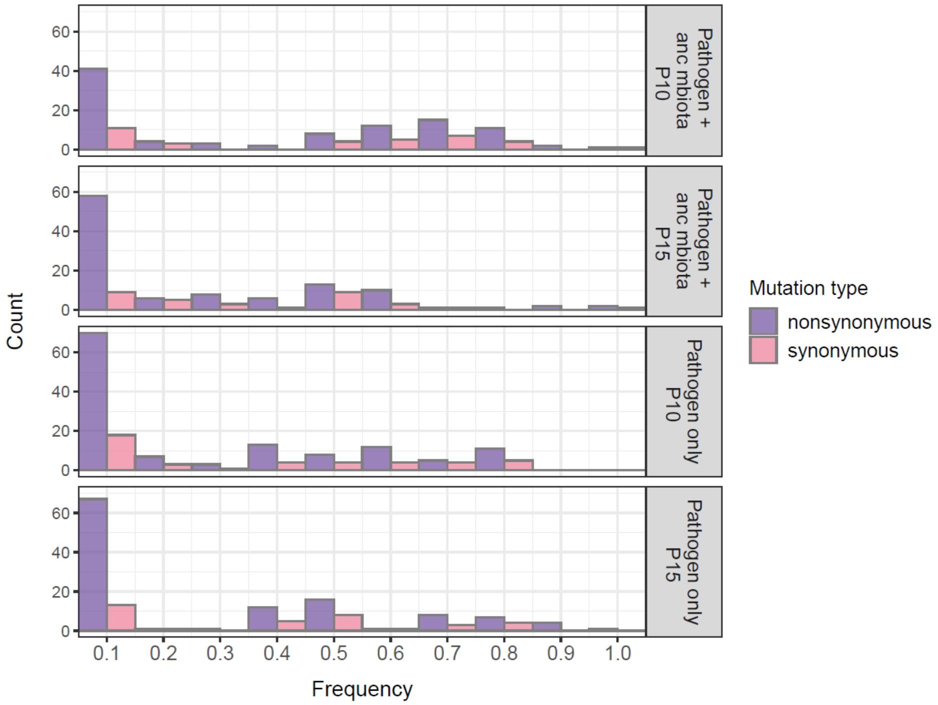
Count
60
40
20
0
Pathogen + anc mbiota P10
Pathogen + anc mbiota P15
Pathogen only P10
Pathogen only P15
0.1
0.2
0.3
0.4
0.5
0.6
0.7
0.8
0.9
1.0
Frequency
Mutation type
nonsynonymous
synonymous

**S5g**

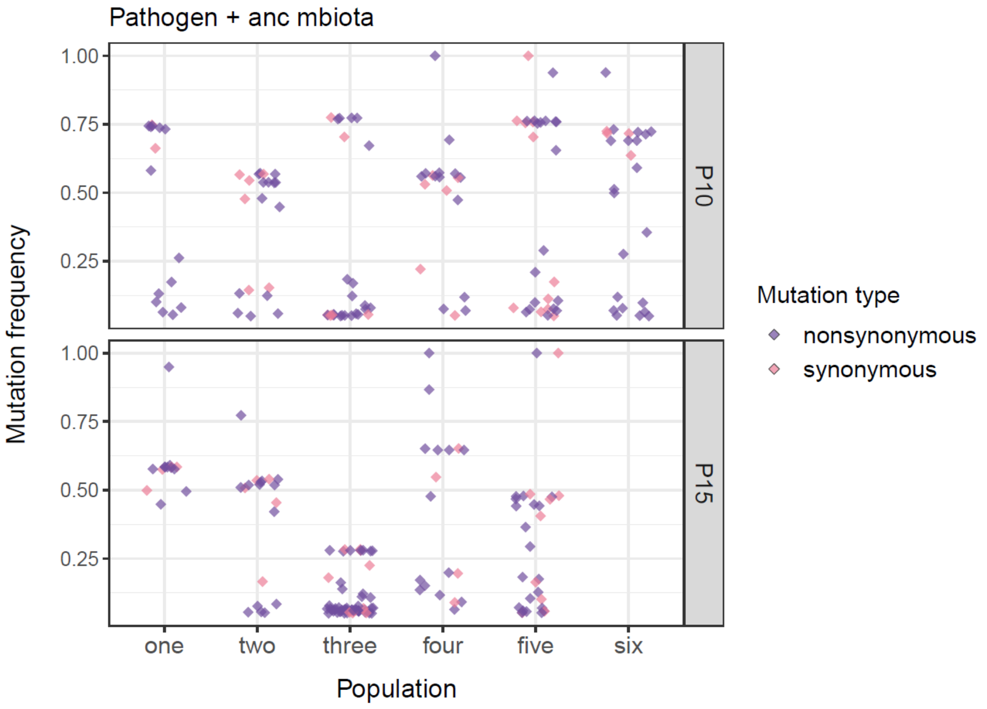

Pathogen + anc mbiota
Mutation frequency
1.00
0.75
0.50
0.25
P10
P15
one
two
three
four
five
six
Population
Mutation type
nonsynonymous
synonymous


**S5h**

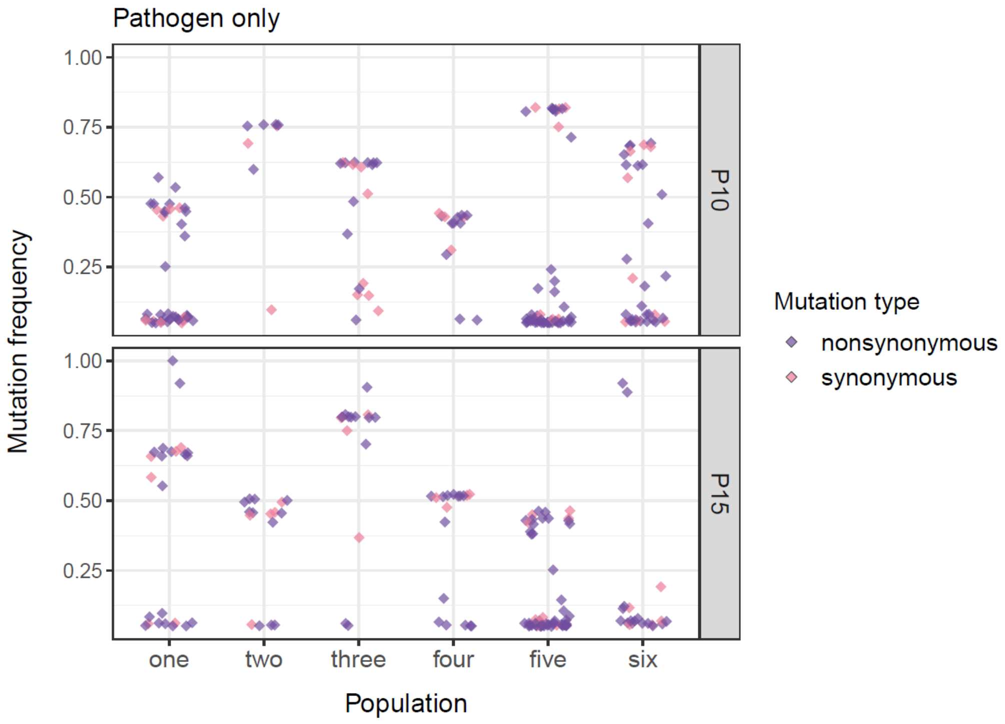

Pathogen only
Mutation frequency
1.00
0.75
0.50
0.25
P10
P15
one
two
three
four
five
six
Population
Mutation type
nonsynonymous
synonymous

**S5i**

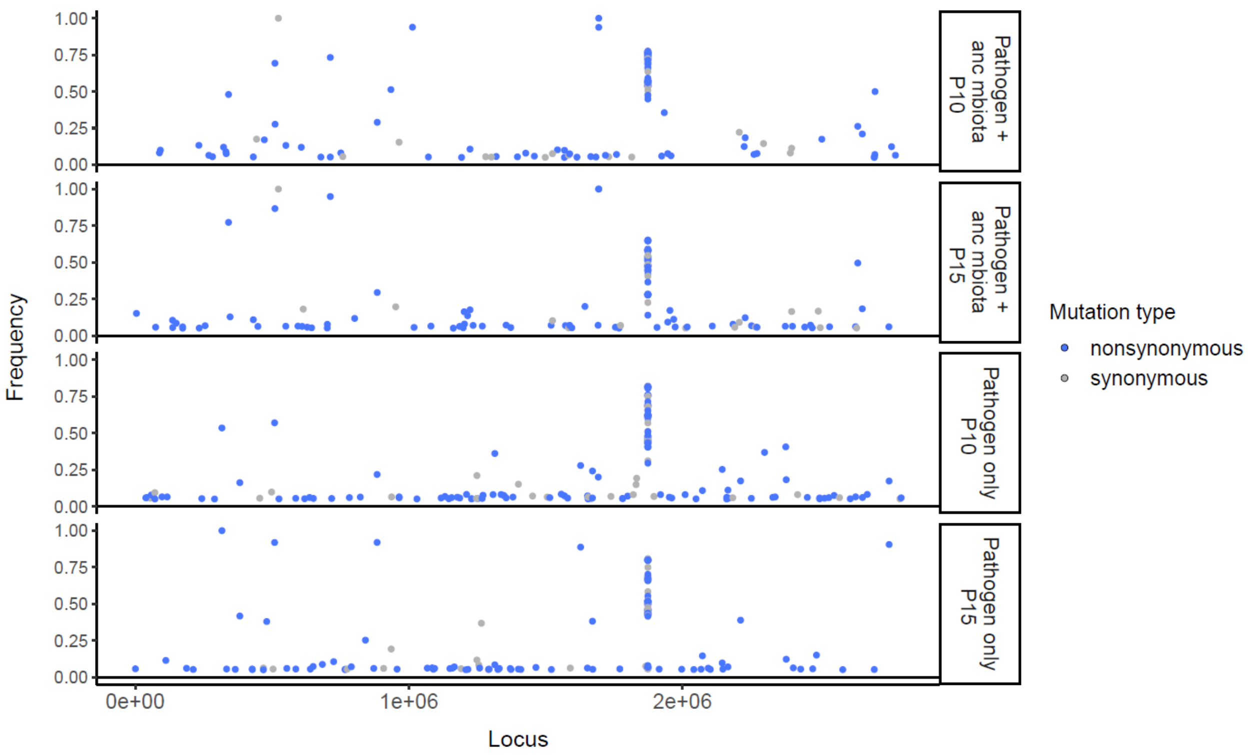

Pathogen + anc mbiota P10
Pathogen + anc mbiota P15
Pathogen only P10
Pathogen only P15
Frequency
1.00
0.75
0.50
0.25
0.00
0e+00
1e+06
2e+06
Locus
Mutation type
nonsynonymous
synonymous

**S5j**

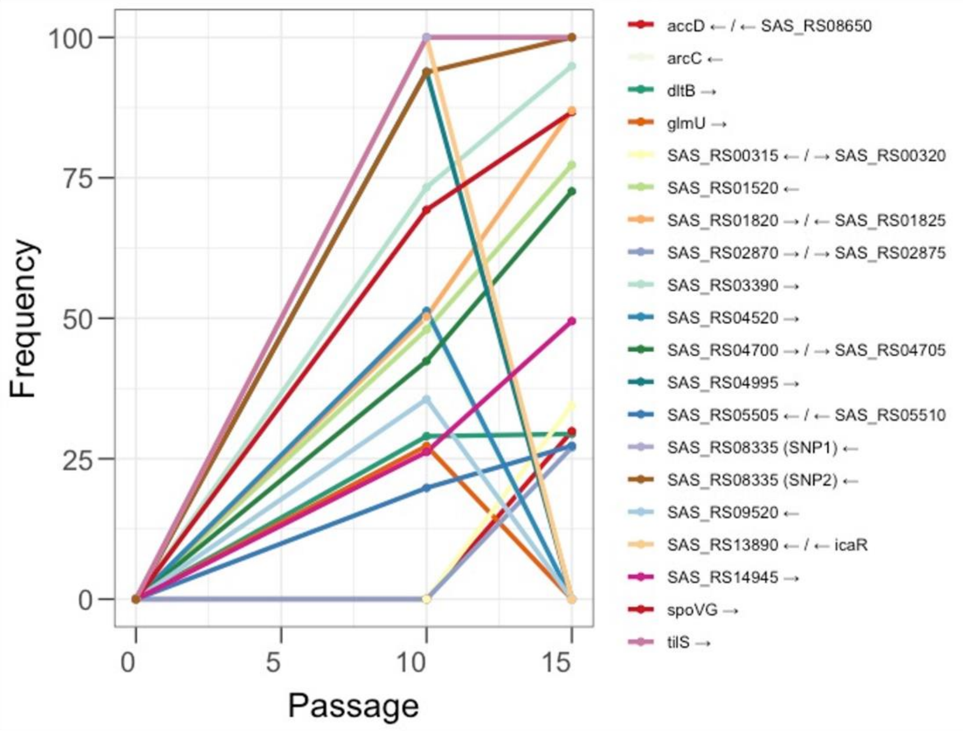

100
75
50
25
0
Frequency
0
5
10
15
Passage
accD ← / ← SAS_RS08650
arcC ←
dltB →
glmU →
SAS_RS00315 ← / → SAS_RS00320
SAS_RS01520 ←
SAS_RS01820 → / ← SAS_RS01825
SAS_RS02870 → / → SAS_RS02875
SAS_RS03390 →
SAS_RS04520 →
SAS_RS04700 → / → SAS_RS04705
SAS_RS04995 →
SAS_RS05505 ← / ← SAS_RS05510
SAS_RS08335 (SNP1) ←
SAS_RS08335 (SNP2) ←
SAS_RS09520 ←
SAS_RS13890 ← / ← icaR
SAS_RS14945 →
spoVG →
tilS →


**S5k**

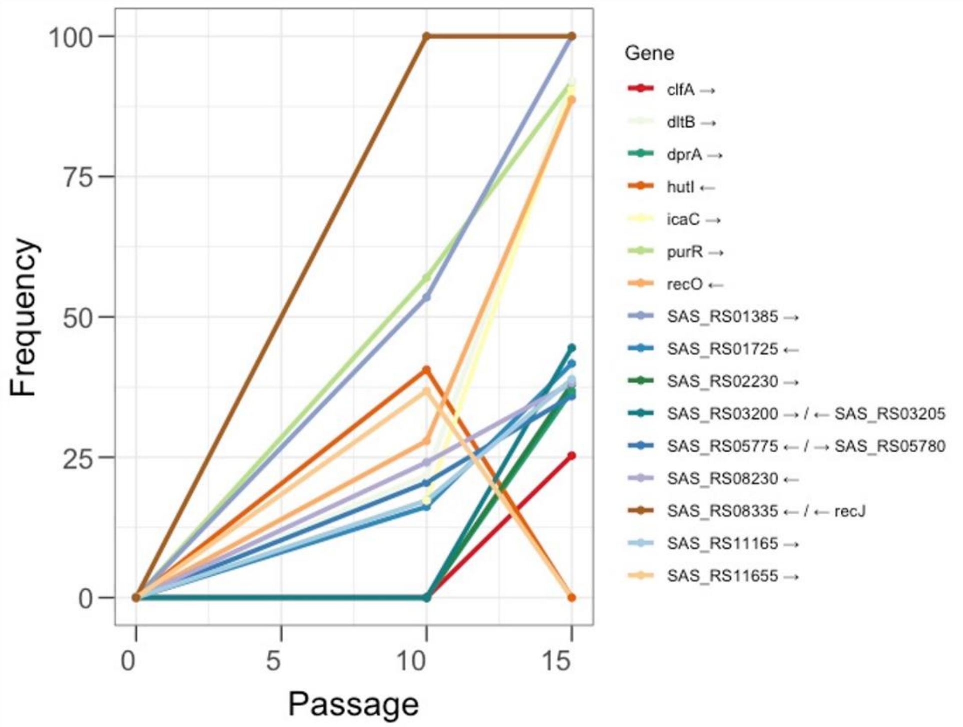

100
75
50
25
0
Frequency
0
5
10
15
Passage
Gene
clfA →
dltB →
dprA →
hutI ←
icaC →
purR →
recO ←
SAS_RS01385 →
SAS_RS01725 ←
SAS_RS02230 →
SAS_RS03200 → / ← SAS_RS03205
SAS_RS05775 ← / → SAS_RS05780
SAS_RS08230 ←
SAS_RS08335 ← / ← recJ
SAS_RS11165 →
SAS_RS11655 →

**S6**

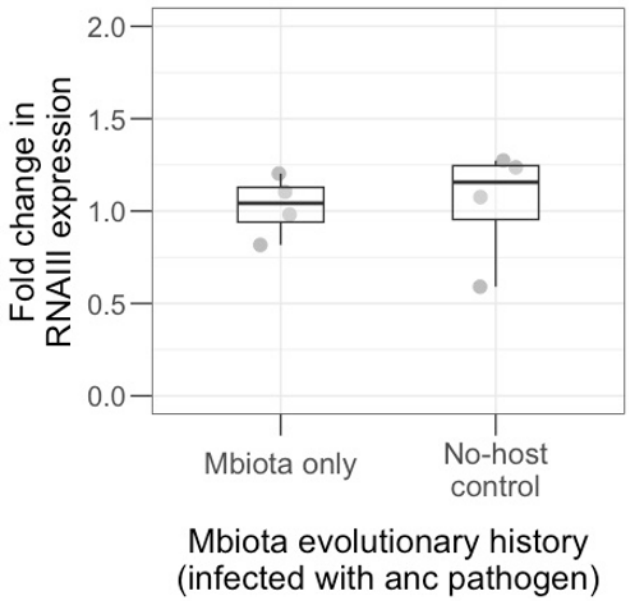
Fold change in RNAIII expression
2.0
1.5
1.0
0.5
0.0
Mbiota only
No-host control
Mbiota evolutionary history (infected with anc pathogen)

**S7**

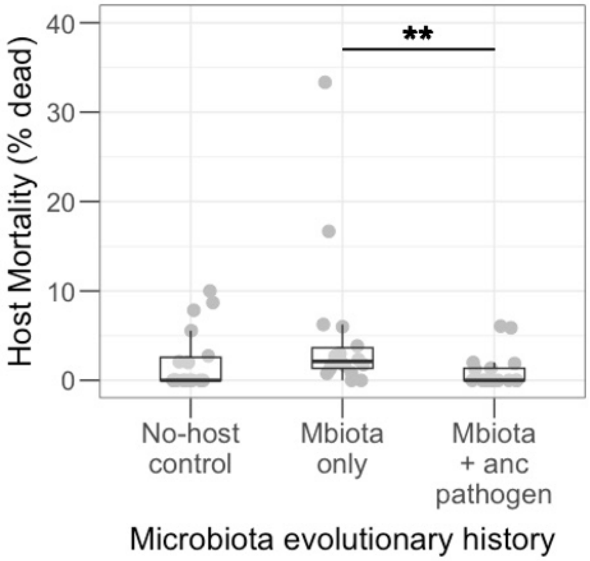
Host Mortality (% dead)
40
30
20
10
0
**
No-host control
Mbiota only
Mbiota + anc pathogen
Microbiota evolutionary history

**S8**

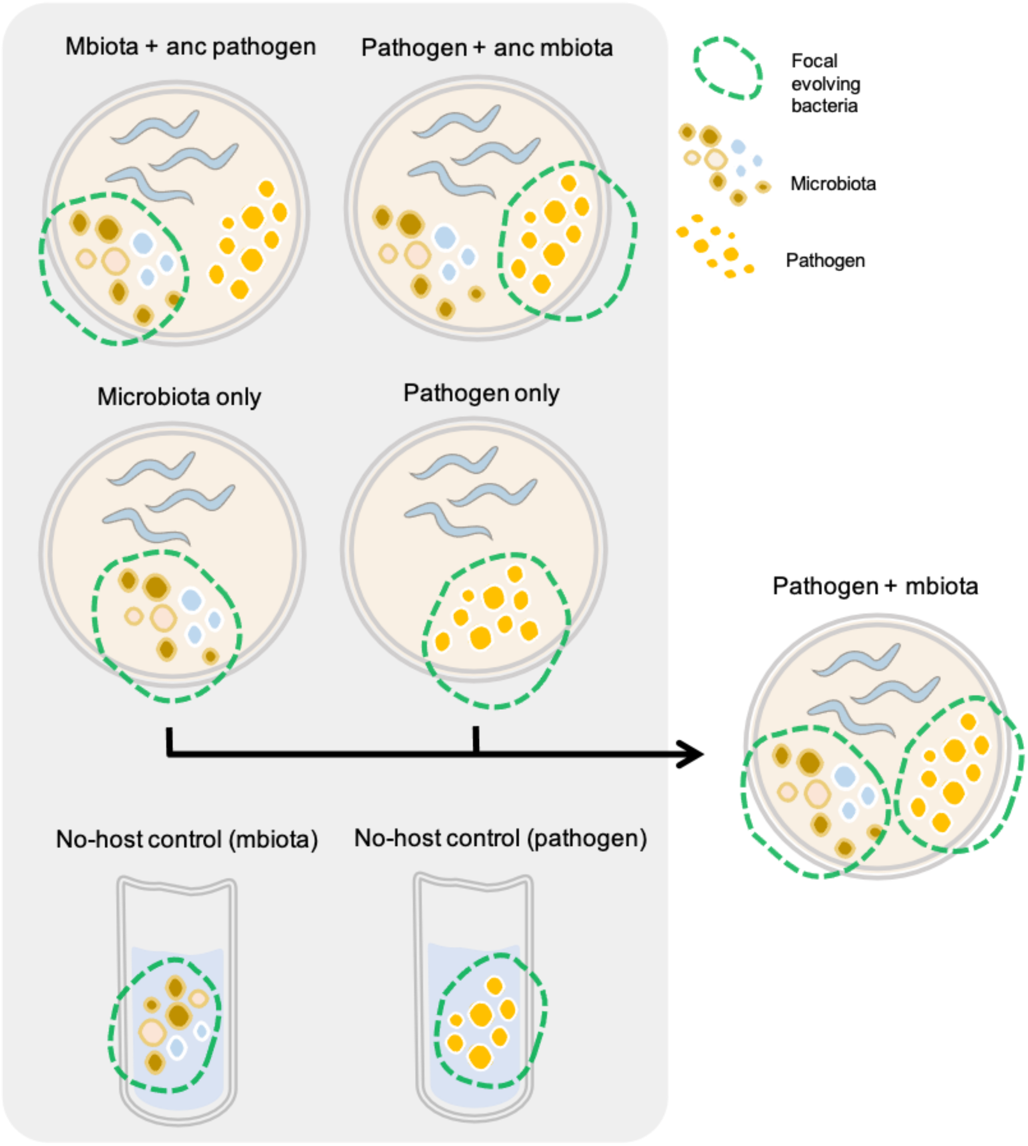
Mbiota + anc pathogen
Pathogen + anc mbiota
Focal evolving bacteria
Microbiota
Pathogen
Microbiota only
Pathogen only
Pathogen + mbiota
No-host control (mbiota)
No-host control (pathogen)

**S9**

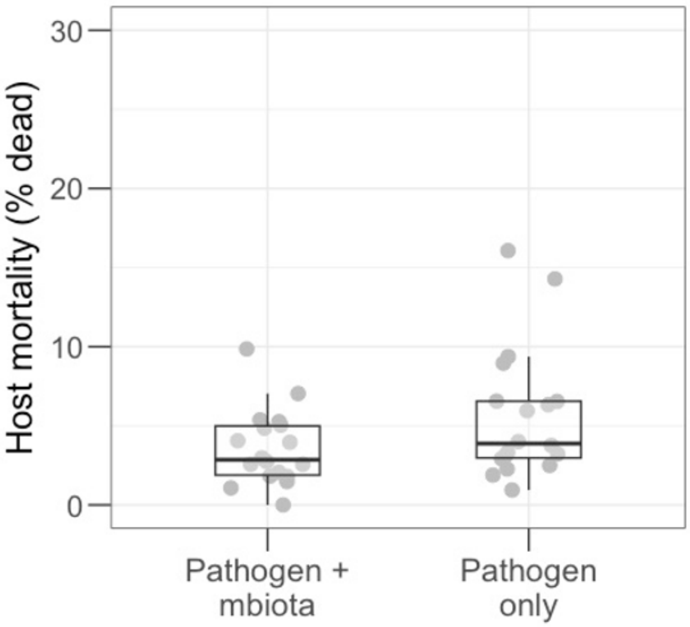
30
20
10
0
Host mortality (% dead)
Pathogen + mbiota
Pathogen only
